# Podoplanin Positive Cell-derived Extracellular Vesicles Contribute to Cardiac Amyloidosis After Myocardial Infarction

**DOI:** 10.1101/2024.06.28.601297

**Authors:** Maria Cimini, Ulrich H.E. Hansmann, Carolina Gonzalez, Andrew D. Chesney, May M Truongcao, Erhe Gao, Tao Wang, Rajika Roy, Elvira Forte, Vandana Mallaredy, Charan Thej, Ajit Magadum, Darukeshwara Joladarashi, Cindy Benedict, Walter J. Koch, Çağla Tükel, Raj Kishore

**Author notes:** **Correspondence to:** Raj Kishore, PhD.

## Abstract

**Background:** Amyloidosis is a major long-term complication of chronic disease; however, whether it represents one of the complications of post-myocardial infarction (MI) is yet to be fully understood.

**Methods:** Using wild-type and knocked-out MI mouse models and characterizing in vitro the exosomal communication between bone marrow-derived macrophages and activated mesenchymal stromal cells (MSC) isolated after MI, we investigated the mechanism behind Serum Amyloid A 3 (SAA3) protein overproduction in injured hearts.

**Results:** Here, we show that amyloidosis occurs after MI and that amyloid fibers are composed of macrophage-derived SAA3 monomers. SAA3 overproduction in macrophages is triggered by exosomal communication from a subset of activated MSC, which, in response to MI, acquire the expression of a platelet aggregation-inducing type I transmembrane glycoprotein named Podoplanin (PDPN). Cardiac MSC^PDPN+^ communicate with and activate macrophages through their extracellular vesicles or exosomes. Specifically, MSC^PDPN+^ derived exosomes (MSC^PDPN+^ Exosomes) are enriched in SAA3 and exosomal SAA3 protein engages with Toll-like receptor 2 (TRL2) on macrophages, triggering an overproduction and impaired clearance of SAA3 proteins, resulting in aggregation of SAA3 monomers as rigid amyloid deposits in the extracellular space. The onset of amyloid fibers deposition alongside extra-cellular-matrix (ECM) proteins in the ischemic heart exacerbates the rigidity and stiffness of the scar, hindering the contractility of viable myocardium and overall impairing organ function. Using SAA3 and TLR2 deficient mouse models, we show that SAA3 delivered by MSC^PDPN+^ exosomes promotes post-MI amyloidosis. Inhibition of SAA3 aggregation via administration of a retro-inverso D-peptide, specifically designed to bind SAA3 monomers, prevents the deposition of SAA3 amyloid fibrils, positively modulates the scar formation, and improves heart function post-MI.

**Conclusion:** Overall, our findings provide mechanistic insights into post-MI amyloidosis and suggest that SAA3 may be an attractive target for effective scar reversal after ischemic injury and a potential target in multiple diseases characterized by a similar pattern of inflammation and amyloid deposition.

**NOVELTY AND SIGNIFICANCE:** **What is known?**

- Accumulation of rigid amyloid structures in the left ventricular wall impairs ventricle contractility.
- After myocardial infarction cardiac Mesenchymal Stromal Cells (MSC) acquire Podoplanin (PDPN) to better interact with immune cells.
- Amyloid structures can accumulate in the heart after chronic inflammatory conditions.

**What information does this article contribute?**

- Whether accumulation of cumbersome amyloid structures in the ischemic scar impairs left ventricle contractility, and scar reversal after myocardial infarction (MI) has never been investigated.
- The pathophysiological relevance of PDPN acquirement by MSC and the functional role of their secreted exosomes in the context of post-MI cardiac remodeling has not been investigated.
- Amyloid structures are present in the scar after ischemia and are composed of macrophage-derived Serum Amyloid A (SAA) 3 monomers, although mechanisms of SAA3 overproduction is not established.

**SUMMARY OF NOVELTY AND SIGNIFICANCE:** Here, we report that amyloidosis, a secondary phenomenon of an already preexisting and prolonged chronic inflammatory condition, occurs after MI and that amyloid structures are composed of macrophage-derived SAA3 monomers. Frequently studied cardiac amyloidosis are caused by aggregation of immunoglobulin light chains, transthyretin, fibrinogen, and apolipoprotein in a healthy heart as a consequence of systemic chronic inflammation leading to congestive heart failure with various types of arrhythmias and tissue stiffness. Although chronic MI is considered a systemic inflammatory condition, studies regarding the possible accumulation of amyloidogenic proteins after MI and the mechanisms involved in that process are yet to be reported. Here, we show that SAA3 overproduction in macrophages is triggered in a Toll-like Receptor 2 (TLR2)-p38MAP Kinase-dependent manner by exosomal communication from a subset of activated MSC, which, in response to MI, express a platelet aggregation-inducing type I transmembrane glycoprotein named Podoplanin. We provide the full mechanism of this phenomenon in murine models and confirm SAA3 amyloidosis in failing human heart samples. Moreover, we developed a retro-inverso D-peptide therapeutic approach, “DRI-R5S,” specifically designed to bind SAA3 monomers and prevent post-MI aggregation and deposition of SAA3 amyloid fibrils without interfering with the innate immune response.

## INTRODUCTION

Amyloidosis is a heterogeneous group of diseases characterized by the extracellular accumulation of stable and poorly soluble cross-beta-sheet monomeric proteins, which are turned into a pathological fibrillar form due to the overproduction of wild-type protein^1^. Amyloid fibrils are rigid, resistant to proteolytic degradation, toxic, and engage intra- and intermolecular interactions with extracellular matrix (ECM) proteins^2,3^. Deposition of pathological fibrillar amyloids in various tissues and organs leads to immune activation and eventually destruction of tissues with impairment of organ function or even organ failure ^4^. Frequently studied cardiac amyloidosis are caused by aggregation of immunoglobulin light chains, transthyretin, fibrinogen, and apolipoprotein in a healthy heart as a consequence of systemic chronic inflammation^1^ leading to congestive heart failure with various types of arrhythmias and tissue stiffness^4 5^. Chronic myocardial infarction (MI) is considered a systemic inflammatory condition, and it has been shown that there is a correlation between plasma levels of hepatic amyloidogenic proteins released after MI and death^6–13^, but whether amyloidosis can also occur after MI is yet to be fully understood. ECM composition of scar tissue after MI has been largely investigated; although fibronectin and collagen are favorable for new myocyte formation, possible deposition of rigid amyloid fibers can reduce the remodeling of the ischemic area, and the onset of amyloid fibers deposition alongside ECM proteins in the ischemic tissue can exacerbate the symptoms of the primary disease and enhances ECM rigidity and stiffness, impairing the scar contribution to left ventricle function after MI^14,15^. Here, we document that amyloidosis occurs after MI and that amyloid fibers are composed of macrophage-derived Serum Amyloid A3 (SAA3) monomers. SAA3 overproduction in macrophages is triggered by exosomal communication from a subset of activated mesenchymal stromal cells (MSC), which, in response to MI, de-novo acquire the expression of a platelet aggregation-inducing type I transmembrane glycoprotein named Podoplanin (PDPN)^16–18^. PDPN acquisition enables MSC to interact and communicate with recruited and activated immune cells CD11b^high^, which highly express the PDPN receptor named C-type lectin-like receptor 2 CLEC-2^19^. Additionally, cardiac MSC^PDPN+^ communicate with different cell types, including macrophages, through the release of extracellular vesicles (EV), including small EVs termed exosomes. Exosomes range from 30 to 150 nm in size, are considered a major paracrine mediator of cell-to-cell communication^20^, and deliver beneficial or detrimental signals to neighboring and distant cells by exchanging their cargo, including protein, non-coding RNA, and lipid^21^. Existing literature suggests that the cellular microenvironment alters exosomal content and function^22,23^ and that during the myocardial injury and repair, exosomes actively participate in intercellular communication between different cell types^24^. Through proteomic analysis of MSC^PDPN+^ derived exosomes (MSC^PDPN+^ Exosomes), we identify SAA 3 protein as an exclusively and highly enriched component of the exosomal cargo^25^. SAA proteins are the major acute-phase plasma proteins that function in innate immunity and are the protein precursor of AA amyloidosis^26,27,28^. They are synthesized largely by hepatocytes but also by extrahepatic cells like macrophages and mesenchymal progenitor cells under the transcriptional regulation of TNFα, IL1α/β, and IL6^29^. Four isoforms of SAA have been identified (SAA1-4)^30^. SAA1, SAA2, and SAA4 are circulating SAAs synthesized in the liver and transported by HDL^31^; however, SAA3 does not contribute to systemic SAA levels, and is expressed only extra-hepatically^31^, and exerts only localized effects at the site of injury. SAA3 binds to toll-like receptor 2 (TLR2) on the macrophage, initiating an autocrine communication to enhance the self-production of SAA3 and the expression of other cytokines and chemokines^32,33^. During chronic inflammation, the abundance of SAA3 derived from activated macrophages impairs the SAA3 clearance, and saturated lysosomes release SAA3 monomers enriched in β-sheets motifs that, being more stable, are prone to aggregate and accumulate to insoluble amyloid protofilaments in the ECM^2,34–36^. Since SAA3 initiates an autocrine SAA3 overproduction in macrophages and that SAAs are precursors of the most common human systemic amyloid diseases worldwide^37^, we investigated whether exosomal SAA3 additionally triggers SAA3 synthesis in macrophages and if SAA3 accumulates in the ischemic hearts in amyloid fibers hindering the contractility of viable myocardium and impairing organ function. We report that exosomes derived from MSC^PDPN+^ isolated from post-MI hearts and injected in normal uninjured mouse hearts lead to immune cell recruitment, fibrosis, cardiac amyloidosis, and depressed cardiac function. MSC^PDPN+ exosome^ delivered SAA3 protein engages with TRL2 on macrophages, triggering an overproduction and impaired clearance of SAA3 proteins, resulting in aggregation of SAA3 monomers as rigid amyloid deposits in the extracellular space. Using SAA3 and TLR2 deficient mouse models, we demonstrated that SAA3 delivered by MSC^PDPN+^ Exosomes has a local function in promoting post-MI amyloidosis. Building on the molecular dynamic simulation, we designed a retro-inverso (anti-sense) peptide made of D-amino acids (DRI) and showed that it is effective in inhibiting SAA3 oligomer aggregation and preventing amyloidosis^38^. DRIs are small hydrophilic proteins (∼5 ammino-acids), enzymatically protected because they are synthesized with D-ammino acids and are minimally immunogenic and not toxic^39,40^. They are naturally eliminated by the urine and exert a specific binding only with the target protein with minimally off-target affinity^39,40^. These characteristics make DRI peptides excellent pharmacological candidates^41–43^. Inhibition of SAA3 aggregation via administration of a retro-inverso D-peptide, specifically designed to bind SAA3 monomers, prevents the deposition of SAA3 amyloid fibrils, positively modulates the scar formation, and improves heart function post-MI.

## METHODS

### Animals

All animal experiments were conducted according to the NIH Guide for the Care and Use of Laboratory Animals and were approved by the Institutional Animal Care and Use Committee (IACUC) of Temple University. Ten- to 12-weeks-old male and female wild type (WT) (C57BL/6J), male and female global Toll Like Receptor 2 knockout (TLR2KO; Jackson Labs; Tlr2tm1Kir/J)^44,45^ and male and female global Serum Amyloid A3 (SAA3) (Jackson Labs; Del(7Saa3-Saa2)738Lvh/J)^46^ knockout mice (SAA3 KO) were purchased from Jackson Research Laboratory (Bar Harbor, ME).

### Animal surgeries

For our physiological assessment and histological analyses, healthy and uninjured ten- to 12-weeks-old male wild type (WT) (C57BL/6J) mice were injected with 2.5×10^8^ exosomes particles isolated from WT or SAA3 KO-MSC^PDPN+^ or murine cardiac endothelial cells (mCECs) conditioned media, in the left ventricle followed by weekly booster doses of same particle numbers by retro-orbital injections. Ten- to 12-week-old male WT, male and female TLR2 KO, and male and female SAA3 KO mice underwent MI surgery previously described in our papers^17^. Briefly, MI was induced through the permanent ligation of the left ventricular descending coronary artery (LAD). Before MI and exosomal treatments, and at 7 and 30 days after both treatments, all animals were screened for baseline and post-MI echocardiography. Following MI, WT animals only were injected i.p. with 100 nmol/kg of DRI-R5S^14^ peptide dissolved in saline once every other day for 30 days or peptide vehicle as a control (saline). At 3, 7, and 30 days after MI, animals were sacrificed under deep anesthesia, bilateral thoracotomy was performed, and the hearts were removed. Mouse hearts at 3, 7, and 30 days after MI were micro-dissected to isolate ischemic and border zone areas from the remote location and lysate in lysis buffer (Cell Signaling Technology) or quiazol (Qiagen). Mouse hearts injected with exosomes were fixed in 4% paraformaldehyde (PFA) and embedded in OCT instead ischemic mouse hearts 30 days after MI were fixed in 10% Formalin (animals subjected to MI) and embedded in paraffin and processed for histological analysis (described below). Furthermore, WT and SAA3 KO mice were euthanized two days after MI, and heart samples were collected for cell culture, as described below.

### Transthoracic echocardiographic analysis and strain

Echocardiography was performed using the Vevo 2100 imaging system from VisualSonics, as publishe^17^. Briefly, transthoracic two-dimensional echocardiography in mice anesthetized with 2 % isoflurane was served with an 18-38 MHz probe. M-mode echocardiography was carried out in the parasternal long axis in mice to assess heart rate (HR), left ventricular (LV) internal diameter in diastole (LVIDd) and systole (LVIDs), LV anterior and posterior wall thickness in diastole and systole (LVAWd and LVAWs as well as LVPWd and LVAWs, respectively). LV fractional shortening (FS%), ejection fraction (EF%), and end-systolic and diastolic volumes were calculated (ESV and EDV)^17^. Speckle tracking strain analysis was done using b-mode loops with a frame rate of 300 at a stable heart rate. M-mode and B-mode data were acquired at baseline and four weeks post-MI in all animals and groups. Global strain and strain rate were analyzed at baseline and post-MI. For all strain analyses, endocardial deformations of the left ventricle were analyzed. All m-mode data analysis was performed using VevoLAB 5.8.1 software, and strain analyses were performed using the Vevo Strain function of Vevo LAB software. All data acquisition and analysis were done blinded.

### In vivo hemodynamic measurements

In vivo, cardiac hemodynamics was performed as described. Mice were anesthetized with a 2% Avertin, and the right common carotid artery was isolated and cannulated with a 1.4 French micro-manometer (Millar Instruments, Houston, TX). LV pressure and heart rate (HR) were measured by this catheter advanced into the LV cavity, and data was recorded and analyzed on a PowerLab System (AD Instruments Pty Ltd., Mountain View, CA). These parameters, as well as maximal values of the instantaneous first derivative of LV pressure (+dP/dtmax, as a measure of cardiac contractility) and minimum values of the instantaneous first derivative of LV pressure (-dP/dtmin, as a measure of cardiac relaxation), were recorded at baseline and after administration of the β-adrenergic receptor agonist, isoproterenol (0.1, 0.5, 1, 5, and 10 ng)^17^. All data were analyzed using LabChart software.

### Histological assessments and Immunohistochemistry of thin cardiac sections

After harvest, either mouse or human hearts were washed with phosphate-buffered saline (PBS), fixed at least for 48h with 4% PFA or 10% formalin, and embedded in OCT (PFA fixed-sucrose gradient) or paraffin (formalin fixed-ethanol and xylene gradient). Cardiac tissues were cut into 5-4 μm-thick sections. Following PBS washing to remove OCT or deparaffinization and rehydration to remove paraffin, mouse samples were stained with Masson Trichrome staining (Sigma), Congo red staining (Sigma), and after either heat-induced or formic acid^47^ antigen retrieval (water bath for 45 min at 90C in citric buffer pH 6.0 with 1% of tween-20), indirectly immunolabeled with commercially available primary antibodies (listed in Supplemental Table 3) and corresponding fluorophore-conjugated secondary reagents (table) or directly stained for the Thioflavin S (Acros Organics) in green. Nuclei were counterstained with 4’,6-diamidino-2-phenylindole dihydrochloride (DAPI; Sigma-Aldrich). Rehydrated slides were also stained for Serum Amyloid A 1-3 after formic acid (70% formic acid in d.d. water) antigen retrieval (20 minutes at room temperature). Multiple sections from the hearts of at least seven mice for each treatment were examined, and representative treatments and time points are described in the results and the Figure’s legend. Images were acquired either with a Nikon Eclipse Ti fluorescence microscope using 4X, 10X, 20X, and 40X objectives or EVOS 7000 (Thermo Fisher Scientific). Masson Trichrome staining images were acquired with a Nikon stereomicroscope at 1X (scar quantification) and a Nikon Eclipse NiE bright field and fluorescence microscope using 4X, 10X, and 20X in bright field. Congo red images were acquired using a Nikon E800 microscope and a digital color camera (Jenoptik Graphax® Kapella) linked to an image analysis system (Bioquant, Life Science 2022 software version, Nashville, TN). Before the use of this microscope for polarized light visualization, a polarized filter was inserted into the appropriate upper slot of the microscope (above the revolving nose piece and below the observation tube), and a lower polarized filter that rotates 360° was attached below the stage (both by Nikon). In addition, the condenser was rotated to the correct differential interference contrast setting for the objective used (DIC-M for 20x and DIC-H for 40X objective, respectively). The Bioquant software’s gain was very dark, and the gain of the red and green colors was set high. The rotation of the lower polarizing filter was slowly moved to approximately 90° until the collagen fibrils showed birefringence colors. Optical sections were projected into a single plane for each color channel and merged using Adobe Photoshop (Adobe) software. Quantitative image analysis was performed with NIH ImageJ by scoring blindly multiple imaging fields.

### Cell isolation and culture

Infarcted WT and SAA3 KO mice were euthanized two days after MI surgery, as described above. The hearts were excised and extensively washed in phosphate-buffered saline. The cardiac tissues were minced and subjected to repetitive rounds of enzymatic digestion with collagenase type 2 (Worthington Biochemical Corp.) until complete dissociation. Larger cells, such as mature myocytes, were precipitated (centrifuged at 100g for 1 minute), and the supernatants containing small cell populations were filtered through 40 µm cell strainers. For positive cell separation, the small cell population was incubated with a biotinylated Podoplanin antibody (table) previously bound with magnetic beads (Milteniy) for 20 minutes at 4C. Cells were sorted through magnetic cell separation columns (Milteniy). After washing, the cells were cultured in an expansion medium on collagen-coated flasks. The purity of the isolation was analyzed by flow cytometry for Podoplanin, LYVE-1 (table), and CD68 (table) on detached (0.2% trypsin) and fixed (4% PFA) cultured MSC^PDPN+^ cells. Lymphatic endothelial cells (LECs) were isolated from WT uninjured mouse hearts with the same methodology. mCECs were purchased from ATCC and cultured expanded. After expansion, all types of cells were cultured in media supplemented with exosome-free FBS and treated with 50ng/mL of TNFα (R&D) and 50nM Angiotensin II (Sigma). Conditioned media was harvested every two days and combined to isolate cell-derived exosomes from the conditioned media (described below). Bone marrow (BM) monocytes were isolated from WT and TLR2 KO BMs mononuclear cells of mice fore and limbs with density-gradient centrifugation as described previousl^17^. Mononuclear cells were seeded on plastic dishes, and the medium was changed after 40 minutes. The monocyte population was cultured and derived in macrophages with 20% FBS, 1% Penicillin/Streptomycin solution, and 20% of L929 conditioned medium in RPMI medium as previously described. Bone marrow-derived macrophages (BMDM), when treated with MSC^PDPN+^ Exosomes, were cultured with expansion media made with RPMI, 20% exosome-free FBS, 1% Penicillin/Streptomycin and 10ng/mL monocyte colony-stimulating factor (Sigma). Wild-type BMDM (∼250,000) were treated with either 10μg, 25μg, or 50μg of recombinant SAA3 (rSAA3) or 1×10^4^, 1×10^5^, 5×10^5^ particles /mL of MSC^PDPN+^ Exosomes for 6, 12 and 24h (Figure S3A). q-PCR analysis for the TLR2 downstream signature cytokines was performed to evaluate the best time point and lowest dose of rSAA3 and exosomal particles needed to activate BMDM. The migratory activity of BMDM was evaluated with a transwell migration assay. BMDM were seeded on the inner side of transwell insets (Corning, 0.8 µm) and let migrated from basal to apical side of transwell insets in different conditions for 24h. BMDM migrated to the apical side were detached with 0.2% trypsin, and the absolute number of migrating cells was counted.

### Single-cell sequencing

Single-cell RNA sequencing of MI hearts was performed as part of a previous study^48^. using the 10× Chromium platform with v 2 Chemistry and analyzed using Cell Ranger v3 (as in https://www.ahajournals.org/doi/full/10.1161/CIRCULATIONAHA.120.044581?rfr_dat=cr_pub++0pubmed&url_ver=Z39.88-2003&rfr_id=ori%3Arid%3Acrossref.org)^48^. Count Matrixes were processed in R version 4.3.1 (https://www.r-project.org/), using Seurat software package version 5.0, by filtering out objects with less than 200 and more than 5000 genes (nFeature_RNA) and a percent. mt over 25. Cell cycle regression, normalization, and scaling were performed on individual objects before merging them following the procedure described in the vignette: https://satijalab.org/seurat/v3.0/merge_vignette.html. The following parameters were used for the clustering: reduction = “pca”, dims = 1:33, resolution =0.5.

### Exosome isolation and characterization

Exosomes were isolated as described in our previous publications^22,24,49,50^. Briefly, 0.2μm filtered conditioned media was concentrated in Amicon ultra centrifugal filter conical tubes from Millipore with a cut-off of 100K, with multiple centrifugations at the maximum centrifugal speed. Exosomes were then isolated with density-gradient ultracentrifugation, and exosome size was analyzed with a nanoparticle tracking analyzer (Nanosight NS300) and expression of exosomal marker proteins verified via Exo-Check™ Antibody Array (System Biosciences, Palo Alto, CA).

### Preparation of cell and tissue lysates and immunoblotting analysis

Exosomes, cellular, and cardiac samples were homogenized using cell signaling technologies 1X lysis buffer (Cell Signaling technologies) enriched with phosphatases and protease inhibitors (Thermo Scientific). Lysates were then centrifuged for 10 min at 4°C at 13,000 rpm to discard the insoluble debris. Next, total protein amounts were quantified via a dye-binding Pierce BCA protein assay kit (Thermo Scientific) and detected using a spectrophotometer reader (SpectraMax i3x Multi-mode Microplate Reader, Molecular Devices) at a wavelength of 512 nm. Equal yields of protein (20-40 μg) were separated through SDS-PAGE and identified by western blot analysis. Total lysates were used to evaluate the exosomal, cellular, and cardiac protein levels. Protein bands were detected by using Odyssey® CLx Imaging System according to the manufacturer’s instructions and quantified with Image Studio™ Lite Software.

### Mass spectrometry analysis

Mass spectrometry analysis was performed by the CHOP-Children Hospital of Philadelphia core facility. Proteins were extracted from exosome samples, denatured, reduced, alkylated, and digested into peptides. They were analyzed in a single shot on QE-HF in DIA mode in the LC-MS/MS analysis system. A spectral library was generated by MaxQuant (DDA), and proteins were quantified using Spectronaut software (DIA, Biognosys). Perseus was used for statistical analysis. Perseus transformed the data using log2, normalized the data by subtracting the median, and using a t-test to identify up/downregulated peptides/proteins between different samples. Biological function and process analysis was performed using gene ontology and KEGG software.

### ELISA analysis

RayBiotech performed mouse cytokine and chemokine 92-plex discovery assay on cell culture supernatant of WT and TLR2 KO BMDM treated with different conditions. Cytokines profile was measured using a 92-plex luminex-based platform. SAA3 levels on MSC^PDPN+^ and mCECs were performed by Eve Technologies (Eve Technologies, Calgary, Canada)^17^ on cell lysates using a Millipore ELISA kit. The SAA3 profile was measured using a 32-plex luminex-based platform. Based on the specific kit used by Eve Technologies, we performed ELISA quantification in-house using the same kit (Millipore) to quantify secreted SAA3 levels on BMDM-conditioned media after treatment with different conditions. SAA3 levels were detected using a spectrophotometer reader (SpectraMax i3x Multi-mode Microplate Reader, Molecular Devices) at a wavelength of 400 nm. The analysis was performed on three different biological samples for each condition.

### Quantitative real-time PCR

Expression levels of different genes were measured using quantitative real-time (RT) PCR technology. Total cellular RNA was isolated from mouse cultured cells and mouse myocardial tissue (scar area and remote area separately) at 3, 7, and 30 days after MI, using miRNeasy Mini Kit (Qiagen) following the manufacturer’s instructions^17^. The cDNA was obtained from total RNA using the high-capacity cDNA revere transcription kit (Applied Biosystems). RT–PCR was performed on an Applied Biosystems 770 apparatus.

### Computational analysis

Using long molecular dynamics simulations, it was proposed in Ref. 60 that the peptide DRI-R5S (sequence: D-SFFSR), built from D-amino acids, inhibits the formation of SAA aggregates and de-stabilizes existing ones. For this peptide, we determined binding sites and the docking for human SAA1 cross-beta fibril structures and mouse SAA3, Fibronectin, and Collagen Iα using the HADDOCK^51^ software. The PRODIGY^52^ software approximated binding energies. We find that the DRI-RSFFS segment interacts with the following mouse SAA3 Monomer Residues: E27, A28, G31, S32, M35, W36, Y39, M42, K43, D51, H55, A62, G68, G69, A72, A73, I76, R80, V83, Q84, T87, H89. The DRI-RSFFS segment disrupts the monomer’s interactions between helix 1 and 3 in its native conformation. A trimmed structure was used to model the effect of a cleaved peptide, similar to human SAA1 (1-76) fragments known to form fibril structures. The structure of the mouse SAA3 monomer was generated from the full mouse SAA3 monomer structure (PDB-ID: 6PXZ) by removing residues 90-122. The guanidinium group of the arginine residue of the DRI peptide nestles into a pocket in the surface of the monomer and contributes most to the interactions between the peptide and monomer (interacting with E27, A28, G31, S32, G68, G69, A72, A73). It should be noted that the DRI peptide interacts with E27 through its backbone atoms rather than its sidechain carboxylate group. The DRI-RSFFS segment interacts with the following Fibronectin Residues (PDB-ID: 1FBR): F5, D6, H7, L21, P22, Y23, M27, V29, G39, I41, T42, C43, T44, S45, R48, C49, N50, R83, G84, E85, W86. These interactions can be organized into three main clusters depending on the DRI residue: Cluster 1 (DRI-residue 1), Cluster 2 (DRI-residues 2-4), and Cluster 3 (DRI-residue 5). Cluster 1 (DRI-residue 1): F5, D6, H7, G39, I4, Cluster 2 (DRI-residues 2-4): L21, P22, Y23, M27, V29, I41, T42, C43, T44, S45, Cluster 3 (DRI-residue 5): T44, S45, R48, C49, N50, R83, G84, E85, W86. The Fibronectin residues of Cluster 3 form a pocket with which the C-terminal arginine residue of the DRI segment interacts. These interactions contribute to most of the contacts observed. Similarly, cluster 1 also forms a pocket on the protein’s surface, interacting with the N-terminus of the DRI-peptide. Cluster 2 contributes to a grove that connects Zone 1 and Zone 3. The final DRI-RSFFS structure was derived from the N-terminal residues RSFFS of human SAA1(hSAA1), which inhibit SAA fibril formation^108^, by reversing the sequence and replacing the common L-amino acids with their D-amino acid analogs. The resulting DRI-peptides are structurally and functionally close to their L-parents, in our case, small hydrophilic and non-toxic molecules that are easily delivered to the organs. However, unlike its L-parents, DRI-R5S is resistant to proteolytic digestion and has a much longer lifetime. Structural models of mouse SAA3 fibrils or oligomers have not been resolved. However, since the mouse SAA3 protein shares 69% amino acid identity with hSAA1 and that hSAA1 fibril structure has been resolved, we could assume in our mouse experiments that DRI-R5S would bind in a similar way to mouse SAA3 than to hSAA1, and that there are preserved aggregation sites (Figure 6a). Binding sites were determined with HADDOCK^51^ software and the binding energies approximated by PRODIGY^52^. As shown, DRI-R5S docks either with mouse SAA3 (Figure 6a) or human SAA1 (Figure S5c) monomer’s sites and simulations performed with fibrils configuration^53^ (Figure S5d) identified that the free energies of binding from the original docking ranged from −23 (SAA1) to −36.4 (SAA3) kJ/mol with the DRI-R5S which is considered an excellent affinity for a small molecule.

### Recombinant and synthesized proteins

Recombinant SAA3 was purchased from Biomatik and reconstituted in sterile water. DRI-R5S (sequence from N terminus to C terminus is SFFSR) was synthesized by Biomatik with serine at the N terminus and arginine at the C terminus with acetate salt endings to enhance the distribution in the tissues. DRI-R5S was reconstituted in sterile and injectable saline water. SB 203580 compound was purchased at Cayman Chemical Company and reconstituted with DMSO.

### Statistical analysis

All in vitro and in vivo experiments during the study period were performed blindly. Data are expressed as means ± SEM. A two-tailed Student’s t-test determined the statistical significance between the two groups. When comparing multiple groups, data were analyzed using a one-way or two-way analysis of variance (ANOVA) test, followed by Tukey’s post hoc test. Differences between groups were considered statistically significant when p≤0.05. All data were analyzed and graphically represented using GraphPad Prism software version 9 (GraphPad, La Jolla, CA).

## RESULTS

### Exosomes from mesenchymal stromal cells positive for Podoplanin (MSC^PDPN+^ Exosomes) induce cardiac injury in healthy mouse hearts

The MSC^PDPN+^ cell population was identified and characterized by our group using flow cytometry and immunohistochemistry analysis (Supplemental Figure 2A) on the cardiac small cell population after MI^16–18^. The Podoplanin expression on a single cell level at different time points after MI was derived using a publicly available published dataset (single-cell RNA sequencing data). These data further confirmed our observations (Supplemental Figure 1)^48^. To study the exosome-mediated communication of MSC^PDPN+^, we isolated MSC^PDPN+^ from mouse hearts two days after MI^17^ (Supplemental Figure 2B), expanded them in culture for 2-3 weeks, and isolated the exosomes from the conditioned media^24^. Culture conditions and purity of cultured MSC^PDPN+^ were previously published by our group^17^. Since MSC in the heart acquire PDPN only after MI^16–18^ (Supplemental Figures 1-2), we verified whether the culture expansion without any stimuli maintains PDPN expression as a marker of stromal cell activation akin to post-MI hearts. We observed that PDPN expression was diminished by the cells in culture (Supplemental Data Figure 2C). To maintain the activated cell status during culture expansion, cells were cultured in medium supplemented with TNFα (50ng/mL) and Angiotensin II (AngII 50nM/mL) two abundantly expressed inflammatory and neurohormonal molecules in the ischemic hearts that trigger PDPN expression and do not initiate cell differentiation^54–59^, and exosomes were isolated from the conditioned medium using ultracentrifugation as per our published studies^22,24^. Exosomes derived from similarly treated mouse cardiac endothelial cells (mCECs), which do not express PDPN, were used as a control to exclude specific effects of *in vitro* TNFα/AngII treatment. In this manuscript, MSC^PDPN+^ Exosomes and mCECs Exosomes are exosomes isolated from MSC^PDPN+^ or mCECs treated with TNFα and Angiotensin II unless otherwise specified. MSC^PDPN+^ Exosomes were analyzed with the nanoparticle tracking analysis to verify size (100-150 nm) (Supplemental Data Figure 2D) and expression of exosomal marker proteins via exosomes markers antibody arrays (Supplemental Data Figure 2E). To understand the effects of the paracrine signals of MSC^PDPN+^ via their exosomes, 2.5×10^8^ exosomal particles isolated from MSC^PDPN+^ or mCECs conditioned media were injected in the left ventricle of healthy 10–12 weeks old C57BL/6J mouse hearts, followed by weekly booster doses of same particle numbers by retro-orbital injections (Supplemental Data Figure 2F). Mice injected with saline were used as a further control. Echocardiography was performed at the baseline and 30 days after initial treatments to observe the effect of exosomal treatments on heart physiology (Figure 1A and 1B). Compared to the baseline, only the group of animals injected with MSC^PDPN+^ Exosomes showed a reduction in the percent ejection fraction (%EF) (Figure 1B) and percent fractional shortening (%FS) (Figure 1B) with an increase in the end-systolic (Figure 1A and 1B) and diastolic (Figure 1A and 1B) volumes (ESV and EDV) suggesting a repression of cardiac function. Histological analysis showed pericardial and infiltrative fibrosis in animals treated with MSC^PDPN+^ Exosomes (Figure 1C, quantified in 1E, left), which was not observed in the animals treated with mCECs derived exosomes (Figure 1D, and 1E, left). The fibrotic tissue in MSC^PDPN+^ Exosomes treated hearts was characterized by extensive infiltration of CD45+ immune cells (Figure 1C right, quantified in 1e, right), which was not observed in the group treated with mCECs derived exosomes (Figure 1D right, quantified in 1E, right). These *in vivo* studies suggest that during ischemic conditions, MSC intracellular communication enhances scar formation, ECM deposition, and immune cell recruitment, making MSC one of the significant cell types during wound healing.

**Figure 1.**
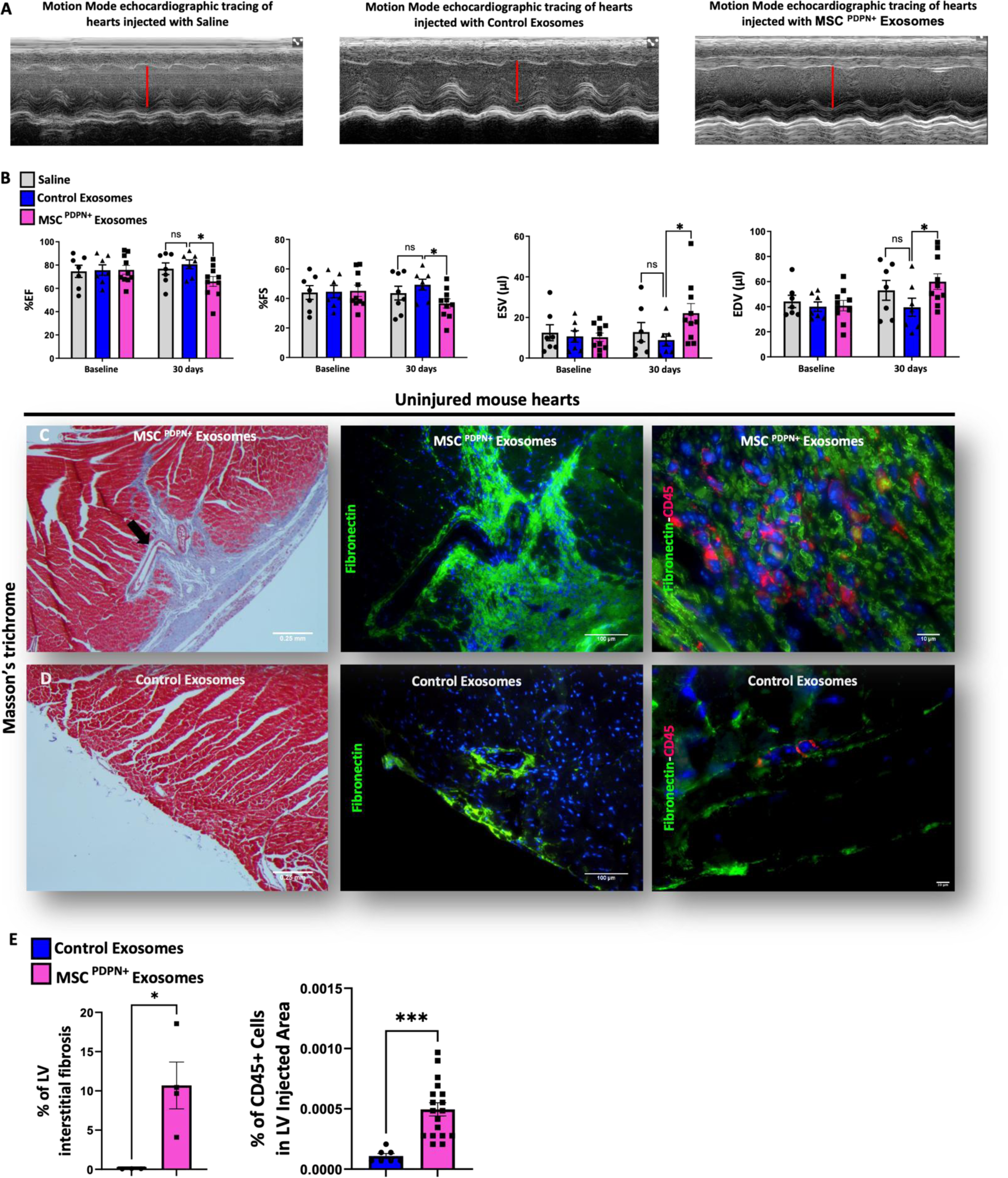
Exosomes derived from Podoplanin positive mesenchymal stromal cells (MSC^PDPN+^ Exosomes) impair cardiac function when injected in healthy mouse hearts. (A) Echocardiographic M-mode analysis of healthy mouse hearts injected with exosomes derived from TNFα and Angiotensin II pretreated MSC^PDPN+^ and mouse cardiac endothelial cells (mCECs) or saline showed dilation of the left ventricle chamber in mouse hearts injected with pretreated MSC^PDPN+^ Exosomes. (B) Echocardiography analysis showed a reduction of ejection fraction (EF) and fractional shorting (FS) percentages and an increase in systolic (ESV) and diastolic (EDV) volumes in healthy mouse hearts injected with pretreated MSC^PDPN+^ Exosomes compared to mouse hearts injected with control exosomes (pretreated mCECs derived exosomes) or saline. Data are presented as mean ± SEM, *P<0,05. N=7-10. Ordinary two-way ANOVA analysis and Tukey’s post hoc test were performed among the groups. (C, left and quantified in E, left). Histological characterization of mouse hearts injected with TNFα and Angiotensin II pretreated MSC^PDPN+^ Exosomes by Masson’s trichrome staining showed infiltrative epicardial fibrosis when compared to animals injected with control exosomes derived from similarly pretreated mouse cardiac endothelial cells-mCECs (D, left). (C, labeled in green, middle, and right panel) Fibrotic tissue was characterized by fibronectin deposition and recruited CD45 positive cells infiltrating the fibrotic tissue (C, right panel, labeled in red and quantified in F, right). Data are presented as mean ± SEM, *P< 0.05, ***P<0.0005. N=7-10. Student’s T-test analysis and Tukey’s post hoc test were performed among the groups.

**Figure 2.**
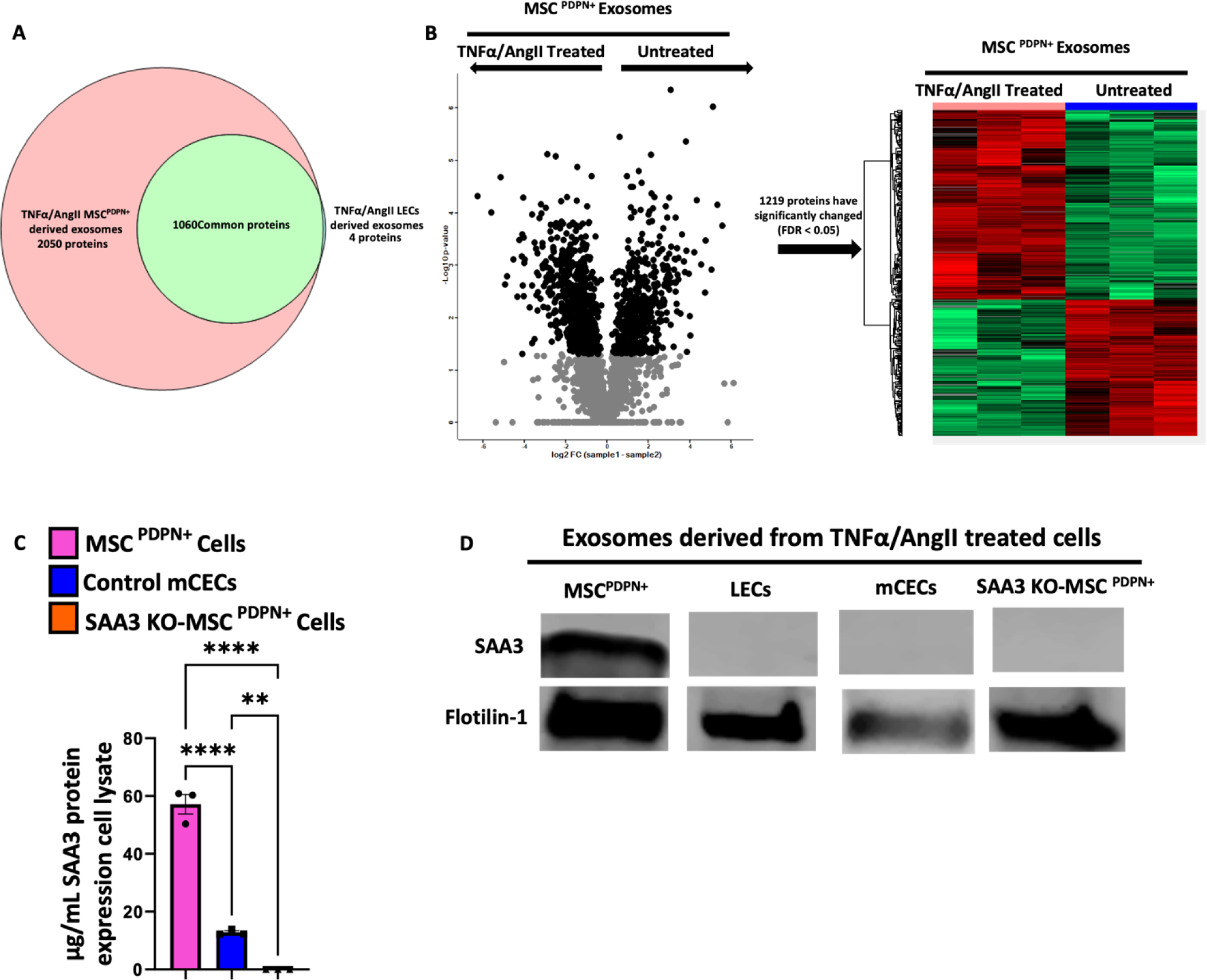
Exosomes derived from Podoplanin positive mesenchymal stromal cells (MSC^PDPN+^ Exosomes) exclusively express Serum Amyloid A3 (SAA3). (A) Mass spectrometry analysis of protein content in exosomes derived from TNFα and Angiotensin II pretreated lymphatic endothelial cells (LECs) and MSC^PDPN+^ cells revealed that ∼1000 proteins are commonly expressed by the two groups of exosomes and that ∼2000 proteins were exclusively expressed in the pretreated MSC^PDPN+^ Exosomes. Within these 2000 proteins, ∼1200 were upregulated after TNFα and Angiotensin II treatment, as shown by the volcano plot (B, left) and the heat map (B, right). (C) The most abundant protein in MSC^PDPN+^ Exosomes is SAA3, which is highly expressed in MSC^PDPN+^’ cell lysate after TNFα and Angiotensin II pretreatment. Treatment with TNFα and Angiotensin II of cardiac endothelial cells (mCECs) slightly increased SAA3 expression in cell lysate, which was undetectable in cell lysate of MSC^PDPN+^ isolated from global SAA3 knockout (KO) mice following the same treatment. (D) Western blot analysis of isolated exosomes derived from TNFα and Angiotensin II pretreated MSC^PDPN+^, LECs, mouse cardiac endothelial cells (mCECs), and SAA3 knockout (KO)-MSC^PDPN+^ showed that SAA3 is solely present in MSC^PDPN^+ Exosomes. Data are presented as mean ± SEM, **P<0,002 and ****P<0.0001. N=3-5. Ordinary one-way ANOVA analysis and Tukey’s post hoc test were performed among the groups.

### Amylogenic Serum Amyloid A 3 (SAA3) protein represents the signature component of mesenchymal stromal cells positive for Podoplanin-derived exosomes (MSC^PDPN+^ Exosomes)

To determine the protein contents within the exosomal cargo that may be responsible for observed inflammation and consequent fibrotic responses in uninjured hearts, we performed proteomic analysis using mass spectrometry on exosomes derived from untreated or TNFα/AngII treated MSC^PDPN+^ or lymphatic endothelial cells (LECs). Since LECs physiologically express PDPN, the goal was to determine whether protein cargo in the exosomes differs between LECs-constitutively expressing PDPN- and MSC^PDPN+^-activating PDPN expression *de novo* in response to MI. LECs were isolated from normal mouse hearts, expanded *in vitro*, and either stimulated or not with TNFα/AngII. Proteomic analysis revealed that treated MSC^PDPN+^ and LECs shared ∼1000 proteins in the exosomal cargo (Figure 2A) and that MSC^PDPN+^ were distinctively enriched in ∼2000 proteins (Figure 2A). Within the exclusive proteins, expression of ∼1200 was significantly changed after TNFα/AngII treatment, as shown by the volcano plot (Figure 2B, left) and the heatmap (Figure 2B, right). Only a few proteins were specifically present either in the untreated or TNFα/AngII treated group (Supplemental Figure 2H). Comparative protein analysis among exclusively expressed proteins in MSC^PDPN+^ Exosomes groups showed that SAA3 was quantitatively and statistically upregulated in MSC^PDPN+^ Exosomes from cells stimulated with TNFα/AngII (Figure 2B). To exclude the effects of TNFα/AngII stimulation, we activated unrelated cells (mCECs) that do not express PDPN and evaluated the impact of the stimulation on SAA3 synthesis. Pretreated mCECs minimally expressed SAA3 in cells after the stimulus (Figure 2C). Similarly stimulated MSC^PDPN+^ isolated from global SAA3 knock-out (KO) mice^46^ (Supplemental Figure 2M) were used as a further negative control and, as expected, did not express SAA3 after treatment (Figure 2C). We confirmed by western blots analysis that SAA3 was exclusively present in exosomes isolated from pretreated MSC^PDPN+^ and was undetectable in exosomes derived from similarly stimulated LECs, mCECs and SAA3 KO-MSC^PDPN+^ (Figure 2D) suggesting that cells naturally expressing PDPN and *de novo* expressing PDPN have a distinct protein profile and the sole expression of PDPN is not a prerequisite for SAA3 expression. Moreover, SAA3 expression in MSC^PDPN+^ Exosomes is cell-specific and not a consequence of *in vitro* TNFα/AngII stimulation, and *de novo* PDPN expression is a prerequisite for SAA3 expression by MSC. Gene ontology analysis of the biological processes (Supplemental Figure 2I), cellular components (Supplemental Figure 2J), molecular functions (Supplemental Figure 2K), and pathways activation (Supplemental Figure 2l) suggested that the exosomal proteins were associated with extracellular matrix modulation, and those proteins associated with exosomes formation indicated that our samples were specifically extracellular vesicles/exosomes (Supplemental Figure 2J). Together, these findings show that SAA3 is exclusively enriched in MSC^PDPN+^ Exosomes^60^, SAA3 expression is a cell-specific characteristic, and only a distinct subset of MSC (MSC^PDPN+^) activated in response to inflammation selectively express SAA3, while PDPN expression (LECs) or proinflammatory stimuli are not sufficient for SAA3 synthesis (LECs and mCECs).

### Exosomal Serum amyloid A 3 (SAA3) is essential to initiate a pathological response in vivo

SAA3 is the major acute-phase protein released at the injury site by immune cells and MSC and functions in innate immunity as both cytokine and chemokine^27–29,61,62^. MSC^PDPN+^ is a heterogenous subset of MSC composed of multiple cell types (PDGFRα^+^, PDGFRβ^+,^ and CD34^+^ cells) that lack common specific markers different from PDPN^16^, whereas PDPN is also expressed by other cell types distinct from MSC like LECs or a sub-category of macrophages^18^ which are the most important source of SAA3^25^, thus, generating a cell-specific knockout mouse model in which only MSC^PDPN+^ does not express SAA3 was impractical. Hence, to evaluate the importance and contribution of exosomal SAA3 in the paracrine communication of MSC^PDPN+^, we injected in healthy mouse hearts MSC^PDPN+^ Exosomes derived from MSC^PDPN+^ isolated from SAA3 KO mice^46^ (Figure 3A). MSC^PDPN+^ from SAA3 KO mouse hearts 2 days after MI were isolated, expanded *in vitro,* and stimulated exactly like wildtype MSC^PDPN+^ before isolation of exosomes from the conditioned media (Figure 3A). Before and after injections of SAA3 KO-MSC^PDPN+^ Exosomes, we followed the animals by echocardiography (Figure 3B), and did not observe any differences in the physiological parameters after exosome treatment, and histologically SAA3 KO-MSC^PDPN+^ derived exosomes induced minimal interstitial fibrosis (Figure 3C, right quantified in d) when compared with healthy mouse hearts injected with wild-type MSC^PDPN+^ Exosomes (Figure 3C, left quantified in D). Our experiment aimed to determine whether SAA3 is one of the essential exosomal proteins involved in the first response of wound healing after injury, and these data provide evidence that MSC^PDPN+^-derived exosomal SAA3 contributes to the observed fibrotic responses and depressed cardiac function seen during the injections of wild-type MSC^PDPN+^Exosomes.

**Figure 3.**
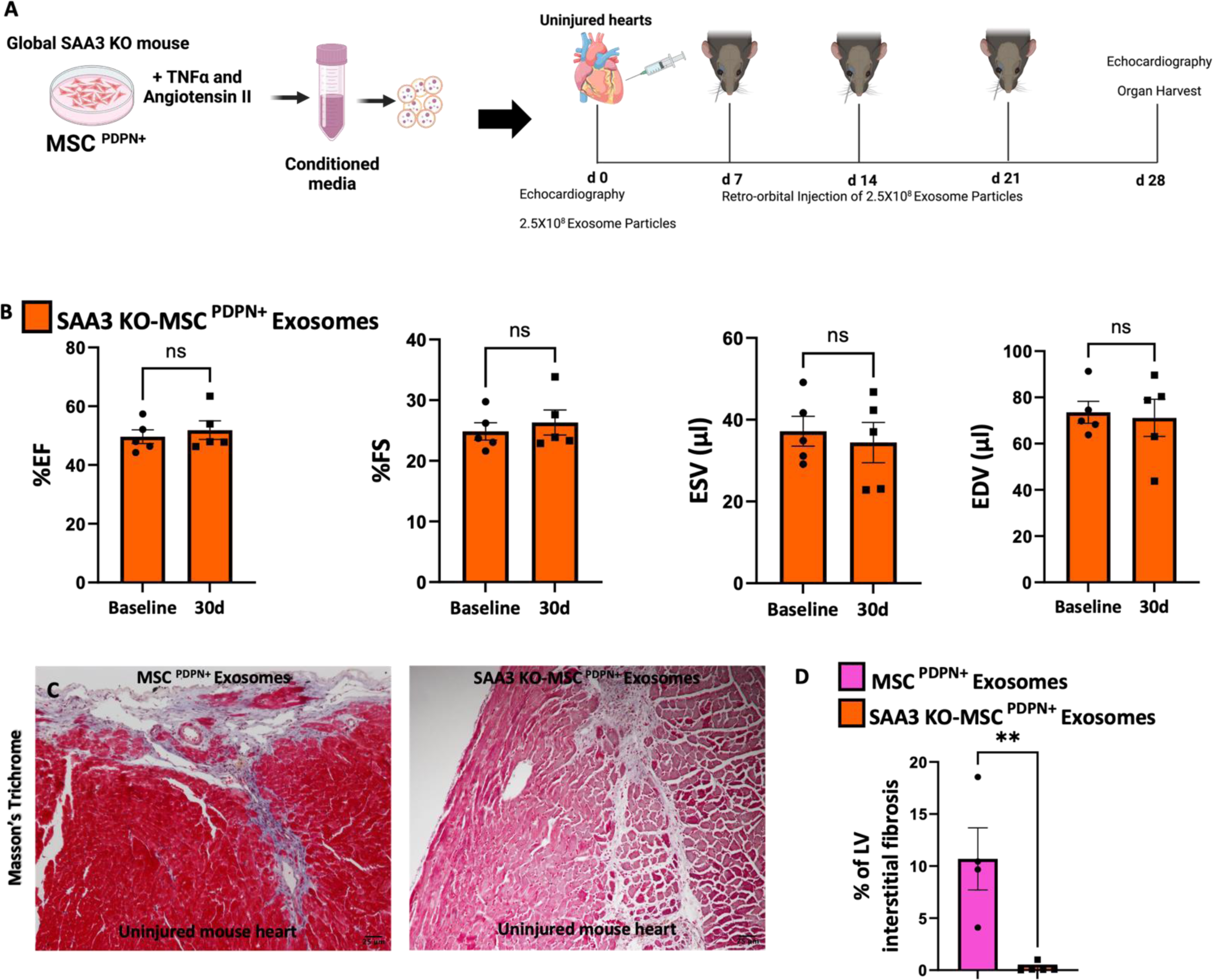
Exosomes derived from Podoplanin positive mesenchymal stromal cells from SAA3 Knockout mice (SAA3-KO-MSC^PDPN+^Exosomes) did not impair cardiac function when injected in healthy mouse hearts. (A) Healthy mouse hearts injected with exosomes isolated from TNFα and Angiotensin II pretreated Podoplanin positive cells (MSC^PDPN+^) isolated from Serum Amyloid 3 knockout mice (SAA3 KO-MSC^PDPN+^) did not show impairment in heart function during echocardiography analysis (B). Data are presented as mean ± SEM. N=5. Ordinary two-way ANOVA analysis and Tukey’s post hoc test were performed among the group. (C, right) Treatment of healthy mouse hearts with exosomes isolated from pretreated SAA3 KO-MSC^PDPN+^ failed to induce epicardial and infiltrative fibrosis compared to exosomes derived from MSC^PDPN+^ treatment (C, left, quantified in D). Data are presented as mean ± SEM, **P< 0.002. N=5-10. Student’s T-test analysis and Tukey’s post hoc test were performed between the groups.

### Exosomal Serum amyloid A 3 (SAA3) and Toll-like receptor 2 (TLR2) are necessary to trigger SAA3 overproduction in macrophages

SAA3 signals through TLR2 and SAA3 can initiate its own overproduction ^32,33^. Thus, we hypothesized that exosomal SAA3 binds TLR2 on macrophages, leading to the activation of downstream pathways and triggering an overproduction of SAA3^63^. To test this, we isolated bone marrow-derived macrophages (BMDM) from wild-type and global TLR2 knock-out (KO)^44,45^ mice, as previously described in our publications^17^ (Supplemental Figure 3A). Wild-type BMDM (∼250,000) were treated with either 1×10^5^ /mL exosomal particles of MSC^PDPN+^ or 10μg of rSAA3. LPS (10ng) and mCECs exosomes (1×10^5^ /mL) were used as controls. 1×10^5^ /mL exosomal particles of MSC^PDPN+^ or 10μg of rSAA3 stimulated the migration of wild-type BMDM from the inner to the apical site of trans well inset (Supplemental Figure 3B)^64^, activated them towards pro-inflammatory phenotype, increasing the expression of proinflammatory cytokines TNFα and IL1β, as well as iNOS2 (Supplemental Figure 3C and 3D) and a wide array of cytokines (Supplemental Figure 3E and 3G) and chemokines (Supplemental Figure 3F and 3H), usually released during innate immunity inflammatory response^64,65^. On the contrary, either MSC^PDPN+^ Exosomes or rSAA3 failed to activate TLR2 KO BMDM (Supplemental Figure 3C-H). Similarly, exosomes derived from SAA3-KO-MSC^PDPN+^ did not trigger the synthesis of hallmark inflammatory mediators in wild-type BMDM (Supplemental Figure 3C, Figure 3E, and 3F). After verifying that MSC^PDPN+^ Exosomes and rSAA3 can initiate a pro-inflammatory phenotype in BMDM, we assessed whether they also triggered SAA3 expression (Supplemental Figure 3A). Wild type and TLR2 KO BMDM were either treated with MSC^PDPN+^ Exosomes, mCECs Exosomes, SAA3 KO-MSC^PDPN+^ Exosomes, or 10ug of rSAA3 for 24h. q-PCR, western blot, and ELISA analysis clearly showed that only wild-type BMDMs were able to express SAA3 (Figure 4A, Figure 4B, and Figure 4C) exclusively after the treatment with MSC^PDPN+^ Exosomes or rSAA3. Exosomes derived from SAA3-KO MSC^PDPN+^ failed to activate TLR2-mediated SAA3 synthesis (Figure 4A, Figure 4B, and Figure 4C). These data show that SAA3 present in MSC^PDPN+^ Exosomes initiate a self-release of SAA3 in macrophages only after binding to TLR2 and that TLR2 is required for SAA3 transcription^66^. Moreover, SAA3 is the required protein present in the MSC^PDPN+^ Exosomes’ cargo that can initiate SAA3 release in competent macrophages. To further corroborate our findings and to dissect the mechanism behind SAA3 synthesis, we verified that the signaling effectors of the TLR2 pathway were activated solely upon binding of rSAA3 with TLR2. Wild-type BMDM were treated *in vitro* with 10μg of rSAA3, and the classical kinases downstream of the TLR2 pathway were analyzed. We did not observe any differences in NFkβ or JNK phosphorylation after 15 minutes of treatment since they are always phosphorylated when monocytes become macrophages; however, treatment with rSAA3 rapidly activated the p38-MAPK pathway, inducing p38 phosphorylation (T180/Y182) (Figure 4D), quantification on the right)^32,46^. Lipopolysaccharide (LPS) was used as a positive control (Supplemental Figure 3I, quantification on the right). We further tested whether SAA3 present in MSC^PDPN+^ Exosomes may independently activate p38 MAPK. We treated BMDM with MSC^PDPN+^ Exosomes and control exosomes. As shown in Figure 4E, 15 minutes of treatment with MSC^PDPN+^ Exosomes resulted in highp38-MAPK phosphorylation compared with baseline untreated wild-type BMDM, which was not observed upon treatment with either mCECs exosomes or SAA3 KO-MSC^PDPN+^ Exosomes (Figure 4E quantification on the right). MSC^PDPN+^ Exosomes treatment, like rSAA3, did not generate differences in the phosphorylation of NFkβ or JNK (data not shown). These data suggested that SAA3 in MSC^PDPN+^ Exosomes engages the p38MAPK signaling pathway upon binding to TLR2. Moreover, treatment of BMDM with 20μM of p38 MAPK inhibitor, SB 203580 compound, decreased SAA3 transcription (Figure 4F) and protein synthesis (Figure 4G, quantification on the right) along with diminished TNFα, IL1β, and iNOS2 expression (Supplemental Figure 3K). LPS was used as a control (Supplemental Figure 3J, quantification on the right). These data show that exosomal SAA3 triggers SAA3 expression in BMDM upon binding with TLR2 and engaging in p38-MAPK phosphorylation (Figure 8).

**Figure 4.**
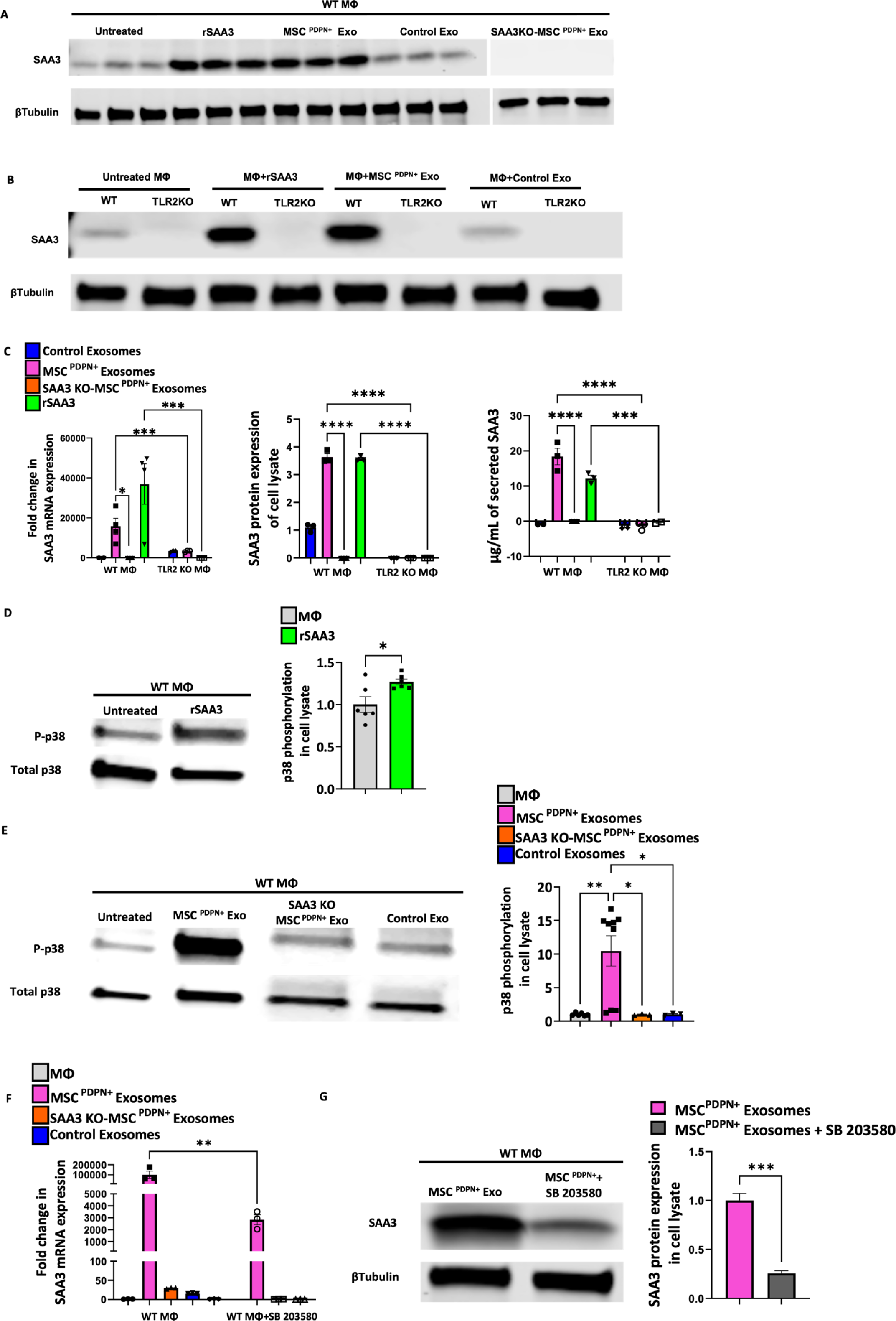
SAA3 over-expression in bone marrow-derived macrophages (MΦ) by MSC^PDPN+^ cell exosomes require Toll-Like Receptor 2 (TLR2) activation and p38-MAPK phosphorylation. (A) Western blot showing of total cell lysate of bone marrow-derived macrophages (MΦ) treated either with recombinant Serum Amyloid 3 (rSAA3), exosomes derived from TNFα and Angiotensin II pretreated Podoplanin positive cells (MSC^PDPN+^ Exosomes), control exosomes derived from similarly treated mouse cardiac endothelial cells (mCECs) or SAA3-null MSC^PDPN+^. MΦ treated with rSAA3 or MSC^PDPN +^Exosomes specifically expressed SAA3 in cell lysates compared to MΦ treated with control exosomes. (B) Western blot showing that MΦ from TLR2 knockout (KO) mice did not synthesize SAA3 upon any stimulation. (C) Quantification of SAA3 expression and release via q-PCR, western blot, and ELISA analysis of wild type and TLR2 KO MΦ after different treatments. Data are presented as mean ± SEM, *P< 0.05, ***P<0,0005, and ****P<0.0001. N=3-5. Ordinary one-way ANOVA analysis and Tukey’s post hoc test were performed among the groups. (D) Western blot analysis and quantification on the right of p38-MAPK phosphorylation (T180/Y182) in bone marrow-derived macrophages (MΦ) after treatment with recombinant Serum Amyloid A 3 (rSAA3). (E) Western blot analysis and quantification on the right of p38-MAPK phosphorylation (T180/Y182) in MΦ after treatment with exosomes isolated from TNFα and Angiotensin II pretreated MSC^PDPN+^. Exosomes from similarly treated MSC^PDPN+^ from SAA3 knockout (KO) mice or murine cardiac endothelial cells (mCECs) did not induce P38-MAPK phosphorylation in bone marrow-derived macrophages (MΦ) (B, quantification on the right). Data are presented as mean ± SEM, *P< 0.05 and **P< 0.002. N=3-6. Student’s T-test analysis, ordinary one-way ANOVA analysis, and Tukey’s post hoc test were performed among the groups. (F) q-PCR analysis of SAA3 mRNA in MΦ shows an important reduction of SAA3 expression after specific inhibition of p38-MAPK with SB 203580 compound. (G) Western blot analysis of SAA3 protein expression shows an important reduction of SAA3 synthesis after specific inhibition of p38-MAPK with SB 203580 compound. Data are presented as mean ± SEM, *P< 0.05 and***P<0,0005. N=3-6. Student’s T-test analysis or ordinary one-way ANOVA analysis and Tukey’s post hoc test were performed among the groups.

### MSC^PDPN+^ Exosomes initiate Serum Amyloid A 3 (SAA3) amyloidosis in healthy mouse hearts

Besides its function during inflammation, SAA3 can be the protein precursor of AA amyloidosis^37^. Based on the literature regarding amyloidosis after MI^6–13,67,68^, we verified with Thioflavin S and Congo red staining, two gold standard diagnostic approaches for amyloidosis^4,69–71^, the presence of amyloid fibers in the scar area of mouse hearts after ischemia. Thioflavin S and Congo red molecules are specific dyes bound by amyloid fibrils, and amyloid deposits bound to Congo red molecules show characteristic birefringence when viewed with a polarizing microscope ^4,69–71^, showing a golden color effect that is required for histological diagnosis^71^. Mouse hearts 30 days after MI showed positivity for Thioflavin S fluorescence (Figure 5A, left panel) and Congo red birefringence (Figure 5B, left panel and Supplemental Figure 4A, left and magnification in the middle) in the ischemic scar tissue (ischemic scar tissue labeled with Masson’s trichrome staining in Supplemental Figure 4B). Positivity to Congo red stain and Thioflavin S confirm the elevated concentration of amyloid assemblies in the cardiac scar. Mechanistically, we showed that macrophages greatly increased the synthesis of SAA3 after triggering the TLR2 downstream pathway. Thus, we confirmed that SAA3 was expressed in the ischemic area of mouse hearts after MI, and the peak of transcription was 3 days after MI, which corresponded to the time of migration and activation of macrophages (Supplemental Figure 4C). We also corroborated the histochemistry data with specific western blot analysis of SAA3 deposition in the ischemic area of infarcted mouse hearts 30 days after MI (Supplemental Figure 4D, quantification on the right). Furthermore, we validated via immunohistochemistry that SAA, labeled in green (Figure 5C, left and 5D, left), aggregates in ischemic tissue of mouse hearts 30 days after MI along with extracellular matrix proteins, labeled with fibronectin in red (Figure 5C, left panel). More interestingly, healthy mouse hearts, without any MI insult, injected with exosomes isolated from TNFα/AngII treated MSC^PDPN+^ showed SAA amyloidosis 30 days after injection (Figure 5A, 5B, and 5C, middle panels, 5B magnification on the right and Supplemental 4E, left). Amyloid deposits were found in the fibrotic epicardial area (Figure 5A, 5B and 5C, middle panels, 5B magnification on the right and Supplemental 4E, left), and Congo red stain showed specific birefringence when observed with polarized light (Figure 5B, middle panel and magnification on the right). Healthy mouse hearts injected with mCECs exosomes did not develop any epicardial amyloidosis (Figure 5A, right panel, and Supplemental Figure 4E, middle). Notably, deposition of SAA was not increased in mouse hearts injected with SAA3-null MSC^PDPN+^ Exosomes (Figure 5C, right panel, and Supplemental Figure 8E, right), and SAA deposits were not observed in the fibrotic tissue or at the site of injection except weak deposition around capillaries (Figure 5C, right panel). This data showed that amyloidosis takes place after MI and that released SAA3 proteins can aggregate in amyloid deposits. The exosomal SAA3, after triggering SAA3 overproduction via TLR2 in macrophages (as described above), represents a potential mechanism of SAA3-mediated cardiac amyloidosis. SAA amyloidosis has never been described in human hearts after ischemic injuries or during fibrotic tissue formation in heart failure^6^. We have previously shown that the cardiac fibrotic tissue of individuals with heart failure is populated by MSC^PDPN+ 17^. We, therefore, verified the presence of SAA amyloid deposits in human failing heart tissue. As expected, infiltrating fibrotic tissue of human heart samples derived from explanted hearts from patients with heart failure (Supplemental Figure 4F) also contained extensive SAA amyloid deposits (labeled in green, Figure 5E, and Supplemental Figure 4G). Taken together, these findings show that exosomes derived from pre-treated mCECs or SAA3-KO MSC^PDPN+^ fail to induce SAA3 amyloidosis in healthy mouse hearts and that SAA amyloidosis can also develop in humans.

**Figure 5.**
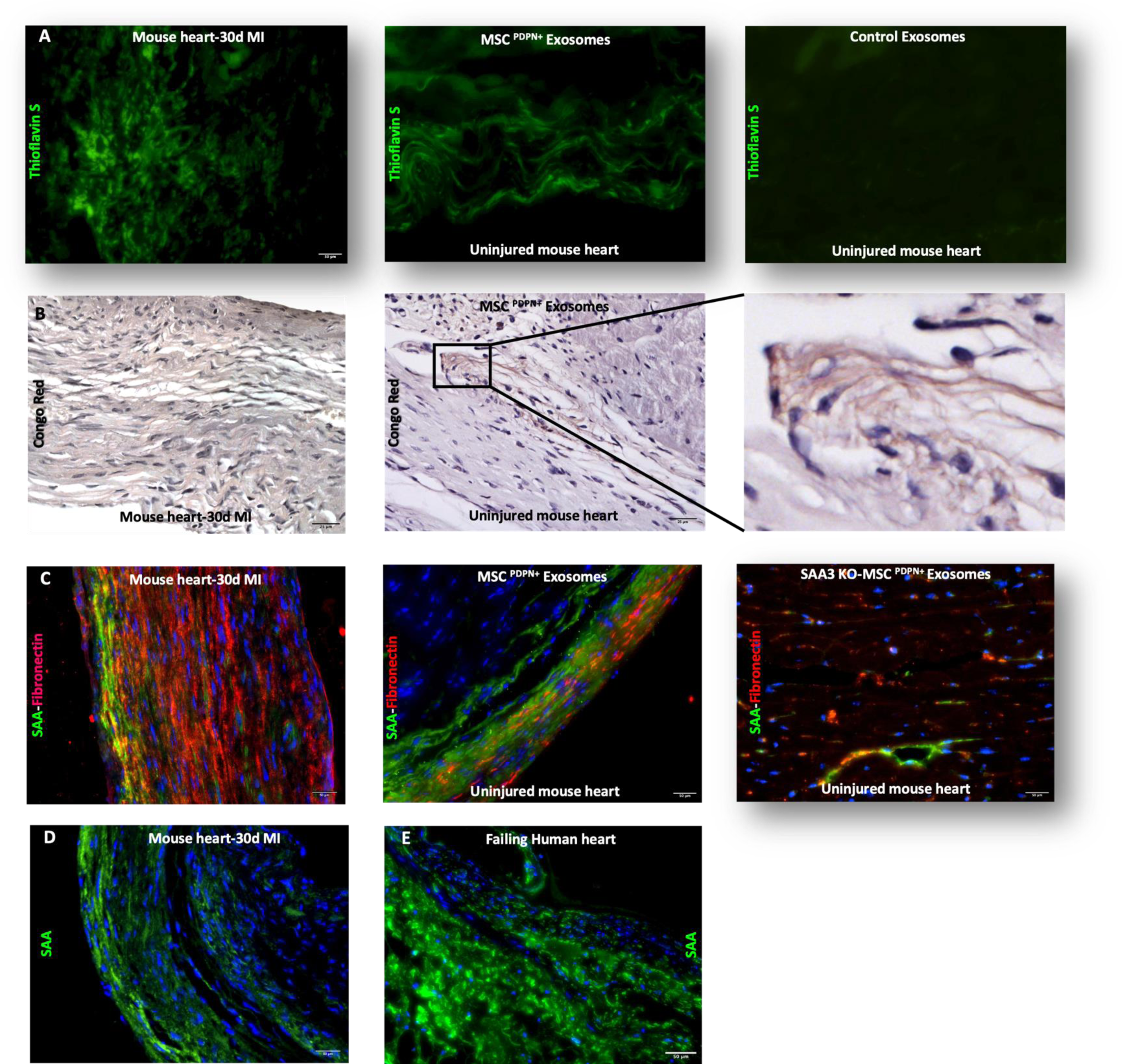
Serum Amyloid 3 (SAA3) aggregates as extracellular amyloid deposits, leading to cardiac amyloidosis. (A, left) Mouse heart sections 30 days after myocardial infarction (MI) and (A middle) healthy hearts injected with TNFα and Angiotensin II pretreated Podoplanin positive mesenchymal stromal cells exosomes (MSC^PDPN+^ Exosomes) were stained with Thioflavin S and Congo red staining (B). Both staining showed the presence of amyloid deposits when compared with healthy hearts injected with similarly treated murine cardiac endothelial cells (mCECs) (a right). Congo red staining also showed specific birefringence (B) of amyloid deposits observed with a polarized light microscope in mouse hearts 30 days after MI or after treatment with MSC^PDPN+^ Exosomes. (C, left) Immunohistochemistry analysis further showed that SAA, labeled in green, aggregated in ischemic tissue of mouse hearts 30 days after MI along with extracellular matrix proteins, labeled with fibronectin in red (C, left), as well as exosomes isolated from pretreated MSC^PDPN+^ were able to initiate a SAA amyloidosis (C, middle) when injected in healthy mouse hearts. (C, right) Conversely, exosomes derived from pretreated MSC^PDPN+^ SAA3-null failed to induce SAA amyloidosis. N=7-10. (B, right) Cardiac sections obtained from failing human hearts showed SAA aggregation, labeled in green, in the fibrotic tissue with the same pattern as SAA aggregation in mouse hearts 30 days after MI (D, left). N=7-10.

### Serum Amyloid A 3 (SAA3) antisense D-Peptide treatment reduces SAA3 oligomerization and amyloidosis after myocardial infarction

As previously mentioned, overproduction of SAA3 impairs SAA3 clearance, and lysosomes release dysfunctional SAA3 monomers that are prone to aggregate to insoluble amyloid protofilaments^2,42^. We measured the aggregation capacity with a protein aggregation assay to validate if SAA3 monomers released by activated BMDM are prone to aggregate in amyloid structure and thereby initiate amyloidosis. Protein aggregation assay^72^ using Thioflavin T showed that after treatment either with MSC^PDPN+^ Exosomes (Supplemental Figure 5A and 5B) or rSAA3 (Supplemental Figure 5A and 5B), BMDM conditioned media is enriched with insoluble amyloid structures that bind Thioflavin T. These data further validated the idea that MSC^PDPN+^ Exosomes and rSAA3 propel a SAA3 feed-forward loop in BMDM and that SAA3 levels are consistently elevated in BMDM conditioned media after rSAA3 or MSC^PDPN+^ Exosomes treatments. Moreover, the presence of SAA3 amyloidosis after MI or injection of MSC^PDPN+^ Exosomes in healthy mouse hearts compelled us to examine whether we can modulate the aggregation of SAA3 protofilaments in vitro and reduce post-MI amyloidosis in vivo. Interestingly, therapeutic approaches targeting sterile innate immunity response have not yet succeeded because impaired inflammatory cell recruitment or neutralization of signature inflammatory cytokine negatively affects the scar formation and healing of the ischemic heart^17,73^. Building on a computational study^38^, we tested DRI-R5S (sequence: D-SFFSR)^38^, an inhibitory peptide that acts as an antisense of the SAA3 binding sites and reduces the aggregation of amyloid fibrils without interfering with the inflammatory response (Figure 6A). The simulations of Jana at al.^38^, showed that the D-Peptide interaction with SAA1-3 monomers made it more difficult for the β-sheets chains to converge into the conformation taken in fibrils. They further showed that DRI-R5S strongly reduced the stability of SAA1 fibrils which makes SAA fibrils growth more difficult (Supplemental Figure 5d). Taking advantage of what has been published on D-peptides as drug candidates^74^ to reduce amyloidosis in Alzheimer’s disease^75–81^, we treated BMDM with 1×10^5^ /mL MSC^PDPN+^ Exosomes along with different doses of DRI-R5S for 24h. We measured the amyloid structures in the conditioned media with Thioflavin T assay (Supplemental Figure 5B). 25μg of DRI-R5S was able to reduce protein aggregation significantly. To further validate the activity of DRI-R5S, we treated BMDM with rSAA3 and 25μg of DRI-R5S for 24h (Figure 6B). DRI-R5S treatment significantly reduced the aggregation activity of amyloid proteins after rSAA3 treatment (Figure 6B). Of note, DRI-R5S did not interfere with rSAA3 binding with TLR2, in fact, after MSC^PDPN+^ Exosomes co-treatment with DRI-R5S, BMDM still expressed SAA3 proteins. Of note, docking analysis of DRI-R5S with fibronectin (Supplemental Figure 5E) and Collagen Iα (Supplemental Figure 5F) proteins showed an affinity for fibronectin and collagen fibrotic structures, signifying that DRI-R5S prevents further interaction of SAA fibrils with ECM proteins. In light of these in vitro results, we treated wild-type animals with 100 nmol/kg of DRI-R5S delivered i.p., every other day for 30 days after the induction of MI (Supplemental Figure 5G). Untreated animals injected only with DRI-R5S vehicle (saline) were used as a control. By Western blot (Figure 6C, quantified in 6D and Supplemental Figure 5H, quantification on the right) and immunohistochemistry (Figure 6E) analysis, we observed a significant reduction of SAA3 and SAA proteins in the scar tissues of mouse hearts treated with DRI-R5S for 30 days after MI surgery. Western blot analysis of micro-dissected scar area of untreated control animals (Figure 6C, quantified in D) showed an increased presence of SAA3 when compared with sham-operated mice. Reduced accumulation of SAA3 in the ECM contributed to better left ventricle functions after ischemia (Figure 7A). Ischemic left ventricles of animals treated with DRI-R5S also showed significant scar reduction (Figure 7C) and viable myocardium composition (Figure 7B, magnification) when compared with the ischemic area of untreated animals (Figure 7B, magnification). The presence of viable cardiomyocytes and possible reduction in scar rigidity and stiffness likely improved the symptoms of the primary ischemia and reduced scar size. To validate our findings and proposed mechanism in wild-type animals, we performed MI in TLR2 global KO and SAA3 global KO animals (Supplemental Figure 6). Wild-type animals injected with saline were used for comparison. After 30 days of MI, both TLR2 and SAA3 KO mice showed an improved heart function as shown by echocardiographic data^5^ (Supplemental Figure 6A), reduced scar size (Supplemental Figure 6C, quantified in D), and increased heart weight/body weight ratio (Supplemental Figure 6E). Ischemic left ventricle wall of KO animals was also characterized by viable myocardium and, importantly, reduction of SAA in the extracellular matrix of TLR2 KO animals (Supplemental Fig 7A, quantified in C). Overall, ischemic tissue of both KO mice and wild-type mice treated with DRI-R5S showed an improved vascularization at 30 days after MI compared to wild-type untreated animals (Supplemental Figure 7B, quantified in D). Functional echocardiographic data were further validated with strain analysis. Strain analysis, the gold standard technique to measure tissue elasticity^14^, didn’t show any differences between % of radial strain in baseline and 30 days after MI in wild-type mice hearts treated with DRI-R5S (Supplemental Figure 6B). This finding indicated an improved synchronous myocardial deformation indicative of improved left ventricular elasticity and, hence, improved left ventricular contractility and stretchiness of the scar tissue (Figure 7B, right). On the opposite, wild-type animals with MI and without any treatment showed a statistical reduction in the % of radial strain (Supplemental Figure 6B), indicating scar stiffness. Both groups of animals at baseline show similar % of radial strain (16.9% and 17.7%). Our findings show that inhibiting SAA3 deposition reduced amyloidosis after MI and improved cardiac functions, reverting scar formation (Figure 8).

**Figure 6.**
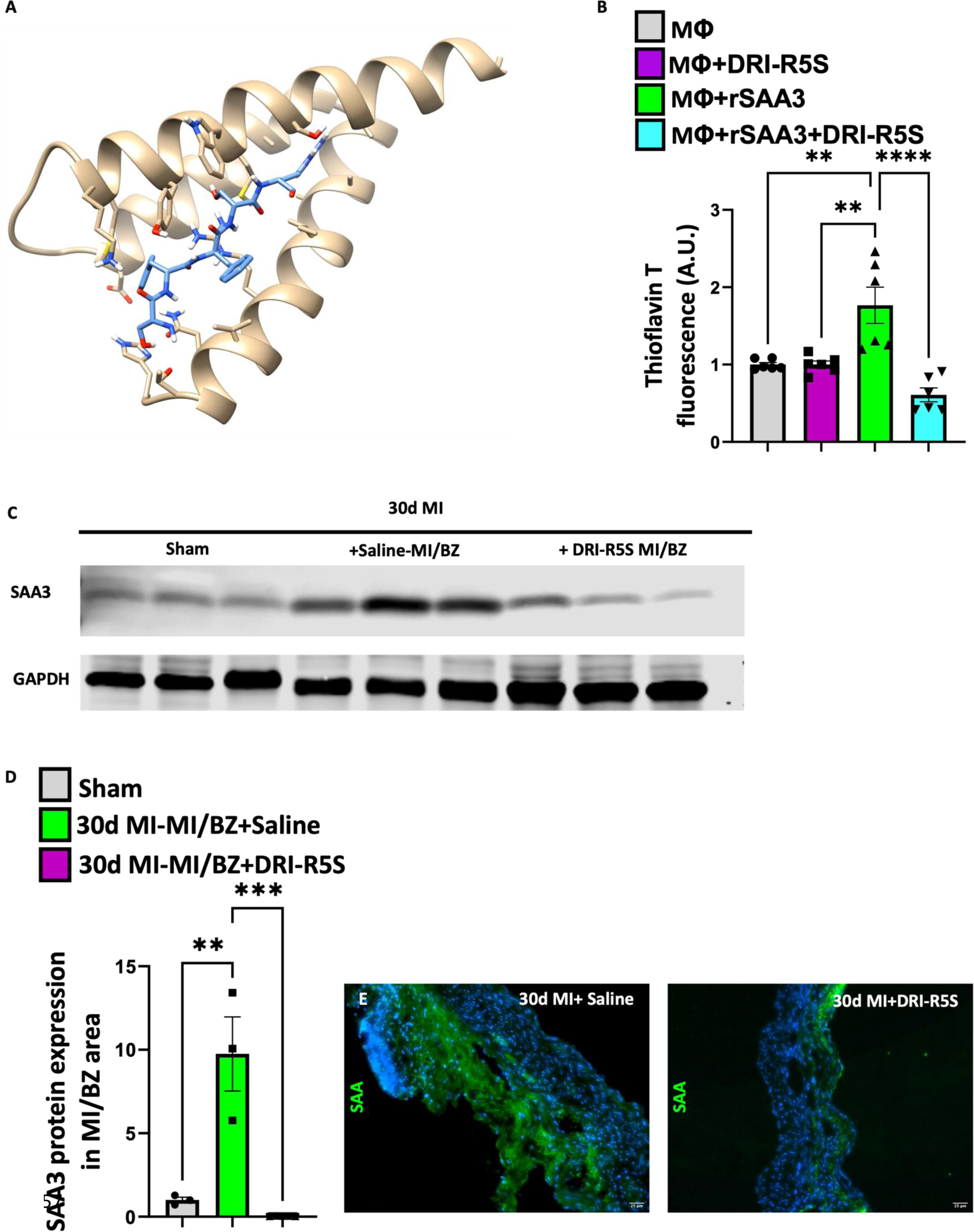
D-Peptide DRI-R5S reduces aggregation of Serum Amyloid A 3 (SAA3) in vivo and in vitro. (A) D-Peptide DRI-R5S docking with mouse SAA3 motif. (B) DRI-R5S reduced the aggregation of SAA3 in vitro in bone marrow-derived macrophages (MΦ) conditioned media after treatment with recombinant SAA3 (rSAA3). (C, quantified in D) In vivo, treatment with DRI-R5S reduced the aggregation of SAA3 in the scar-border zone area of mouse hearts after myocardial infarction (MI). (E) Specific immune-histological staining for SAA showed reduced deposition of SAA in the ischemic area of mouse hearts 30 days after MI when compared with untreated mouse hearts 30 days after MI. Data are presented as mean ± SEM, **P< 0.002, ***P<0.0005 and ****P<0.0001. N=3-5. Ordinary one-way ANOVA analysis and Tukey’s post hoc test were performed among the groups.

**Figure 7.**
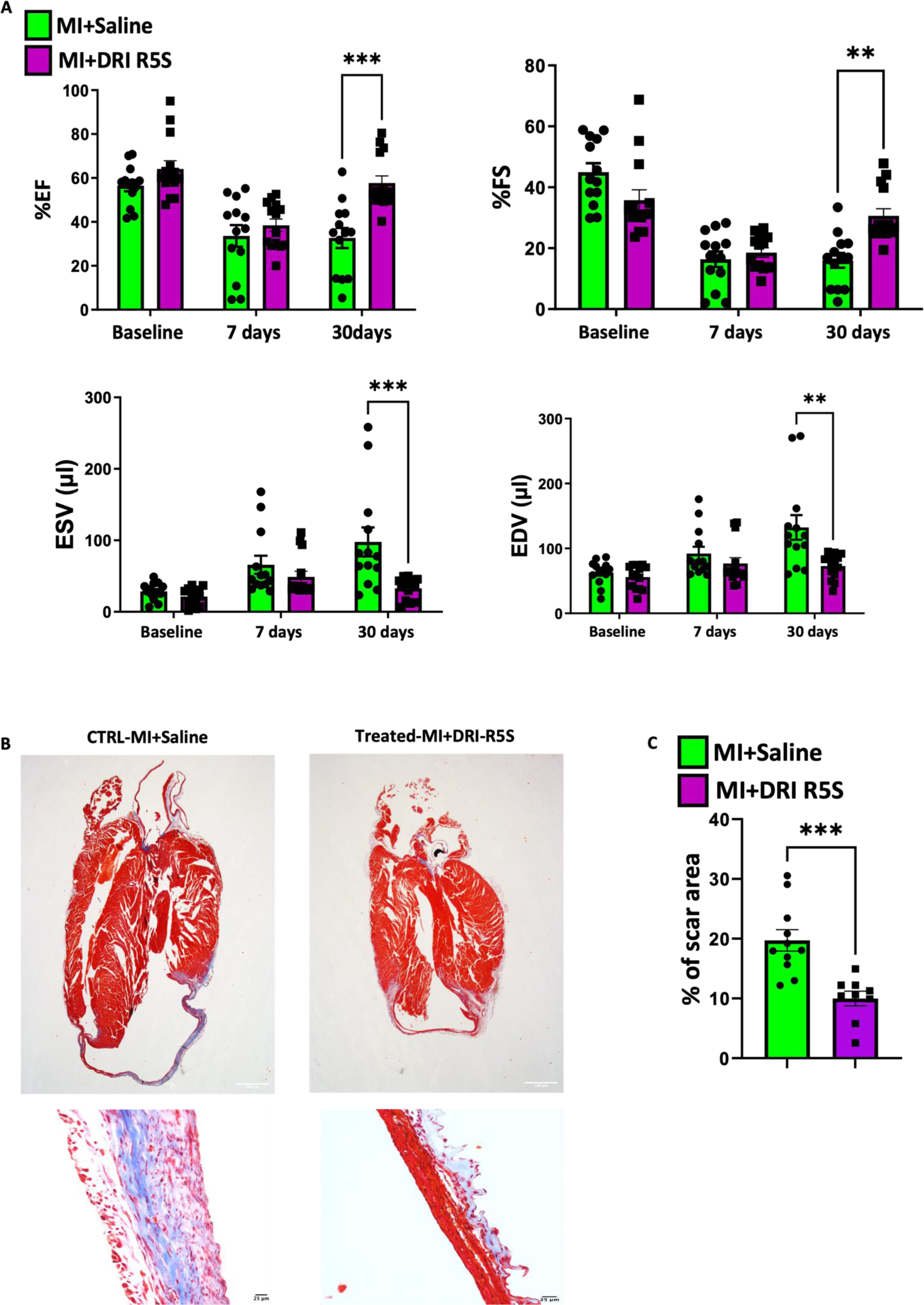
D-Peptide DRI-R5R improves cardiac function after myocardial infarction (MI). (A) Echocardiographic analysis of wild-type animals that underwent MI and were either treated or not with D-Peptide DRI-R5S. Treated animals showed improved cardiac function. (B) Masson’s trichrome staining of wild-type animals that underwent MI and were either treated or not with D-Peptide DRI-R5S showed reduced cardiac scar size (quantified in C) and better-left ventricle wall composition (B, magnification). Data are presented as mean ± SEM, *P< 0.05, **P<0.002, and ***P<0.0005. N=7-10. Ordinary two-way ANOVA (echocardiography data) or student’s T-Test analysis and Tukey’s post hoc test were performed among the groups.

**Figure 8.**
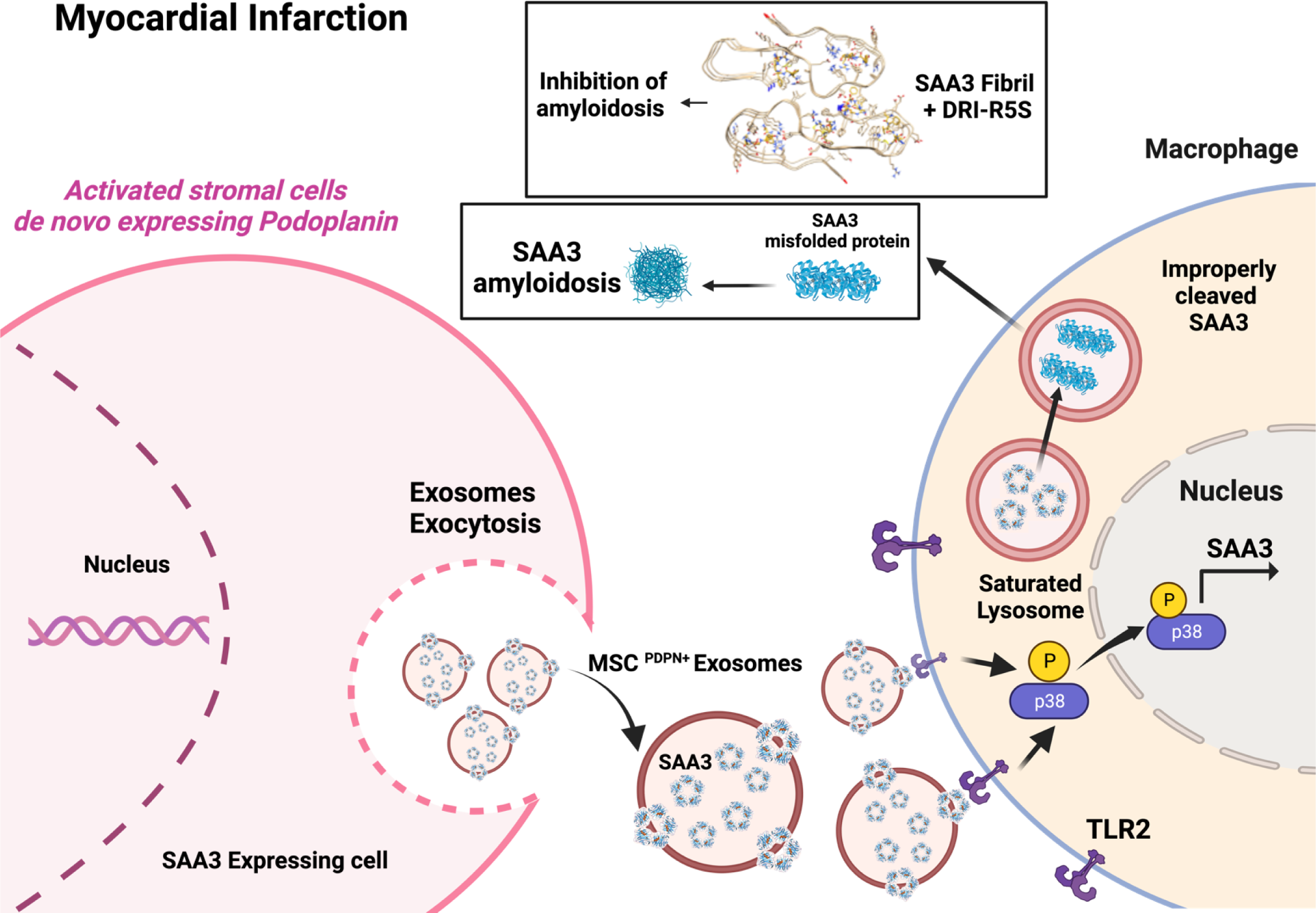
Mechanism and rescue of cardiac amyloidosis after myocardial infarction (MI). Mesenchymal stromal cells positive for Podoplanin (MSC^PDPN+^), continuously stimulated by inflammatory and neurohormonal signals from the microenvironment of the ischemic heart, release Serum Amyloid A 3 (SAA3) through their exosomes (MSC^PDPN+^ Exosomes) prolonging macrophage recruitment and inflammation via engaging with to Toll-like Receptor 2 (TLR2). SAA3 triggers an overproduction of SAA3 in macrophages, with consequent impaired clearance of SAA3. Saturated lysosomes release SAA3 monomers that are prone to aggregate as extracellular, insoluble, and rigid amyloid deposits, leading to cardiac amyloidosis. Inhibition of SAA3 monomers and fibril aggregation using a D-Peptide DRI-R5S reduces amyloidosis after myocardial infarction and improves cardiac functions.

## DISCUSSION

In this study, we described that amyloidosis could take place in the cardiac scar after MI and that it is mainly caused by aggregation of SAA3. SAA3 aggregation is a consequence of exacerbated macrophage activation by MSC^PDPN+^ Exosomes. This mechanism presupposes the binding of exosomal SAA3 with TLR2 and activation of the TLR2 pathway with final phosphorylation of p38. To bypass this process specifically in the injury site, we directly interfered with amyloid fibers formation using a retro-inverso D-Peptide DRI-R5S that binds SAA3 on self-aggregating sites. Our study revealed that DRI-R5S is a powerful pharmacological tool that impaired the onset of amyloidosis during scar formation after ischemia. Reduction of cumbersome structures in the ECM promoted a healthier scar formation after MI, resulting in a muscularized and vascularized left ventricular wall.

### Podoplanin, a possible mesenchymal stromal cells inflammatory marker

Cardiac homeostasis is maintained through well-organized interaction between parenchymal cardiomyocytes and multiple other cell types to exchange signals in both physiological and pathological conditions^82,83^. Cardiac MSC play a distinct role in supporting cardiac growth and structure and in the maintenance of overall tissue homeostasis^84^. After MI, the tissue homeostasis is altered, and the physiological adaptation to a pathological condition consists in activating the highly specialized response cells of wound healing, including immune cells^85–87^. Among all the specialized cells, MSC de novo acquires a glycoprotein named Podoplanin (PDPN)^16,18^ as a sign of activation after ischemia^16,18,48,84^. During embryogenesis, PDPN is essential for the heart and lymphatic system formation^88–93^. In adult life, PDPN is physiologically expressed by Lymphatic Endothelial Cells (LECs) and Fibroblastic Reticular Cells (FRCs) in lymph nodes and allows immune cells to adhere to LECs and FRC to facilitate migration and antigen presentation^94–96^ After MI, PDPN acts as an inflammatory marker and newly acquired PDPN by MSC help them to interact and communicate with recruited and activated immune cells CD11b^high^ which highly express the PDPN receptor named C-type lectin-like receptor 2, CLEC-2^19^. The interaction between PDPN and CLEC-2 activates an inflammatory response in immune cells^97,98^, and we recently showed that blocking the interaction between MSC positive for PDPN (MSC^PDPN+^) and macrophages improved cardiac function and scar composition after infarction^17,97,98^. PDPN acquisition by MSC has been observed and described in various pathologies, including cardiovascular diseases like atherosclerosis, thrombosis, and aneurysms, but also during the progress of autoimmune diseases like psoriatic and rheumatoid arthritis, in the neuropathology of stroke, Alzheimer’s disease, traumatic brain injuries, and a variety of neuroinflammation. Additionally, PDPN expression has been observed during bacterial infections in the gut, inflammatory bowel disease, preeclampsia, and fallopian tubes inflammation and in a large variety of solid tumors. All pathological conditions characterized by PDPN acquisition are listed in Supplemental Table 1. Taking into consideration that cells are described and classified by their markers and that many pathologies are characterized by MSC^PDPN+^, we can conclude that PDPN can be recognized as an established mesenchymal cell marker, specifically and solely expressed during inflammation or pathological conditions. At the moment, there isn’t a consensus on MSC inflammatory markers, and PDPN can be the first of many. Additionally, PDPN is an already established marker for solid tumor prognosis but given its expression in a variety of pathologies, it can be considered a marker for chronic inflammatory conditions as well. In the context of heart failure, not much is known about MSC^PDPN+^, and our studies can add important information about the role of MSC in pathological conditions in addition to the studies that already characterized MSC activity using other different markers as a sign of activation. Our discoveries can open a new research avenue in the study of cardiovascular disease, and the fact that PDPN expression in MSC occurs during or after immune activation raises the possibility that it is a conserved phenomenon that may be valid in all immune reactions and chronic inflammatory diseases. Indeed, looking at the findings on MSC^PDPN+^ in other pathologies, our understanding is that cell-to-cell or paracrine communication between MSC^PDPN+^ and immune cells represents a non-canonical or alternative innate immunity response, and that is an additional trigger for the production and release of proinflammatory cytokines and chemokines including SAA3.

### Innate immunity and cardiac amyloidosis

SAA3 is a powerful pro-inflammatory enhancer of macrophage activation and recruitment (via TLR4), lymphocyte T-helper differentiation in Th-17^+^, and secondary SAA3 amyloidosis^32,33,46,64,99^. We show that SAA3 aggregates in amyloid structures in the scar after ischemia in murine and human hearts. Cardiac amyloidosis is an extensively studied field; prominently investigated cardiac amyloidosis are secondary and caused by aggregation of immunoglobulin light chains, transthyretin, fibrinogen, and apolipoprotein in a healthy heart as a consequence of systemic inflammation^1,2,4,26,42^. Once these proteins create amyloid structures in the heart, the cardiac physiology declines, and the heart goes into failure. However, studies regarding amyloidosis after ischemia, mechanisms behind amyloid fibril accumulation, or which type of amyloid protein may aggregate after MI are yet to be reported. The fact that cardiac amyloidosis occurs as a secondary phenomenon of an already pre-existing chronic condition does not exclude the possibility that it can also occur after prolonged myocardial ischemia (chronic myocardial infarction) or any other types of heart failure, which are considered chronic inflammatory conditions as well^6,7,100^. Our group not only reported the presence of amyloid proteins in the human and murine-failing hearts but also associated the onset of SAA3 amyloidosis with an alternative intracellular communication between macrophages and MSC^PDPN+^ (Figure 8). No prior study has reported a biological and functional role of MSC^PDPN+^ Exosomes in cell-to-cell communication in the ischemic myocardium, which may be relevant due to the sustained presence in the scar tissue (70% of the scar’s cells population) and enhanced communicatory capabilities of activated stromal cells that de novo acquire PDPN^16,18^. Previous studies on chronic cardiovascular diseases had shown correlations between plasma levels of hepatic isoforms of SAA (SAA1 and SAA2) and death. Patients with MI or elevated ST and a high plasma level of SAA1-2 have a higher possibility of dying^6–13,67,68^. Causes of death after high plasma levels of SAA1-2 are unknown. However, SAA1-2 can accumulate in a healthy heart as a secondary amyloid structure, but at the moment, there are no investigations on whether plasma SAA1-2 can also aggregate in the ischemic scar after MI in addition to locally released SAA3. SAA3 is an inflammatory protein that is very conserved and described as one of the major 25 inflammatory markers in a variety of diseases^86^. A literature review regarding pathological conditions specifically characterized by SAA3 overexpression, deposition as amyloidosis, and acquisition by mesenchymal stromal or other cell types is listed in Supplemental Table 2. The type of pathologies in which SAA3 amyloidosis is present and SAA3 is the dominant cytokine unexpectedly result in diseases similar to those in which PDPN is de-no acquired by MSC (Supplemental Table 1). Thus, there may be a correlation between PDPN acquisition by MSC and SAA3 overexpression and consequent amyloidosis. Besides evidence reported in this study, PDPN acquisition and SAA3 amyloidosis were never associated before; although our findings are restricted to cardiac ischemia, based on what is published on SAA3 expression and MSC^PDPN+^, we can predict a similar PDPN expression in MSC, SAA3 overexpression in macrophages, and resultant amyloidosis in other organ systems. Moreover, SAA3 can represent, in addition to PDPN, an in situ inflammatory marker. SAA3 and PDPN are new critical players during inflammation but are involved in mechanisms that cannot be abrogated or inhibited. Blocking TLR2 or interfering with innate immunity activation has been shown in many clinical trials to be detrimental rather than helpful. Thus, we approached the problem using an alternative system that does not interfere with the innate immunity response and is currently under investigation for the treatment of protein-aggregating diseases like Alzheimer’s disease.

### DRI as a pharmacological tool

Retro-inverso D-peptide is an attractive tool with excellent pharmacokinetics, pharmacodynamics, and little off-target effects. These small peptides specifically bind the sequence for which they were designed, and unless they were created to block the interaction between ligands and receptors, they do not interfere with paracrine communications^39–41^. We took advantage of these characteristics and developed a D-Peptide, DRI-R5S, to inhibit SAA3 aggregation^38^, resulting in efficiency and specificity^101,102^. Our peptide does not interfere with the binding of TLR2 with SAA3 or with any other pathway involved in innate immunity; thus, innate immune reaction and inflammation were not inhibited during our treatment^73^. This is fundamental for proper scar formation. Therapies with retro-inverso D-peptides, to our knowledge, have been proposed in the past for the treatment of Alzheimer’s disease, but the design or the employment of D-peptides for the treatment of cardiovascular disease has never been done before. We are the first to suggest using a retro-inverso D-peptide to inhibit protein aggregation after MI as a safe tool to enhance scar reversal without engaging in regeneration mechanisms but only modulating a process that otherwise would have been exacerbated. Our results confirmed that inhibiting the aggregation of cumbersome structures without blocking any physiological reaction is not only propaedeutic for a healthy scar formation, but the treatment is safe and restricted to the timing of macrophage activation and resolution of inflammation, which makes a good point for the translational application of our proposed treatment (Figure 8). Treatments with D-peptides can be beneficial for the treatment of all diseases characterized by SAA3 overexpression/amyloidosis, for example, to avoid the growth of atherosclerotic plaques or to inhibit the aggregation of other kinds of amyloid proteins since, currently, there are no specific treatments for cardiac amyloidosis.

## CONCLUSION

SAA3 amyloidosis occurs after MI and further impairs ischemic heart function. We developed a safe tool that does not inhibit or interfere with the communication between MSC and macrophages but can indirectly modulate the exacerbation of scar formation post-MI, abrogating the onset of SAA3 amyloidosis (Figure 8). Besides MI and heart failure, many pathologies characterized by MSC^PDPN+^ and SAA3 amyloidosis could benefit from our findings. Moreover, this new treatment may work alongside existing therapies.

## ACKNOWLEDGMENTS

M.C. conceptualized this study, designed and performed experiments, analyzed data, and wrote and revised the manuscript. U.H.E.H. designed and performed computational experiments and analyzed DRI-R5S data. C.G. performed experiments. A.D.C. performed computational experiments and analyzed DRI-R5S data. M.M.T. performed experiments. E.G. performed myocardial infarctions and conducted a blinded hemodynamic analysis. T.W. performed myocardial infarctions. R.R. conducted blinded echocardiography and analyzed the data. E.F. analyzed single cell sequencing data. V.M. performed experiments. C.T. performed experiments. A.M. performed experiments. D.J. performed experiments. C.B. performed experiments. W.J. K. edited manuscript. Ç.T. helped in conceptualizing this study and designing experiments. R.K. conceptualized the study, designed experiments, and finalized and approved the manuscript. All authors have given approval to the final version of the manuscript.

## SUPPLEMENTAL MATERIAL

**Supplemental Figure 1.**
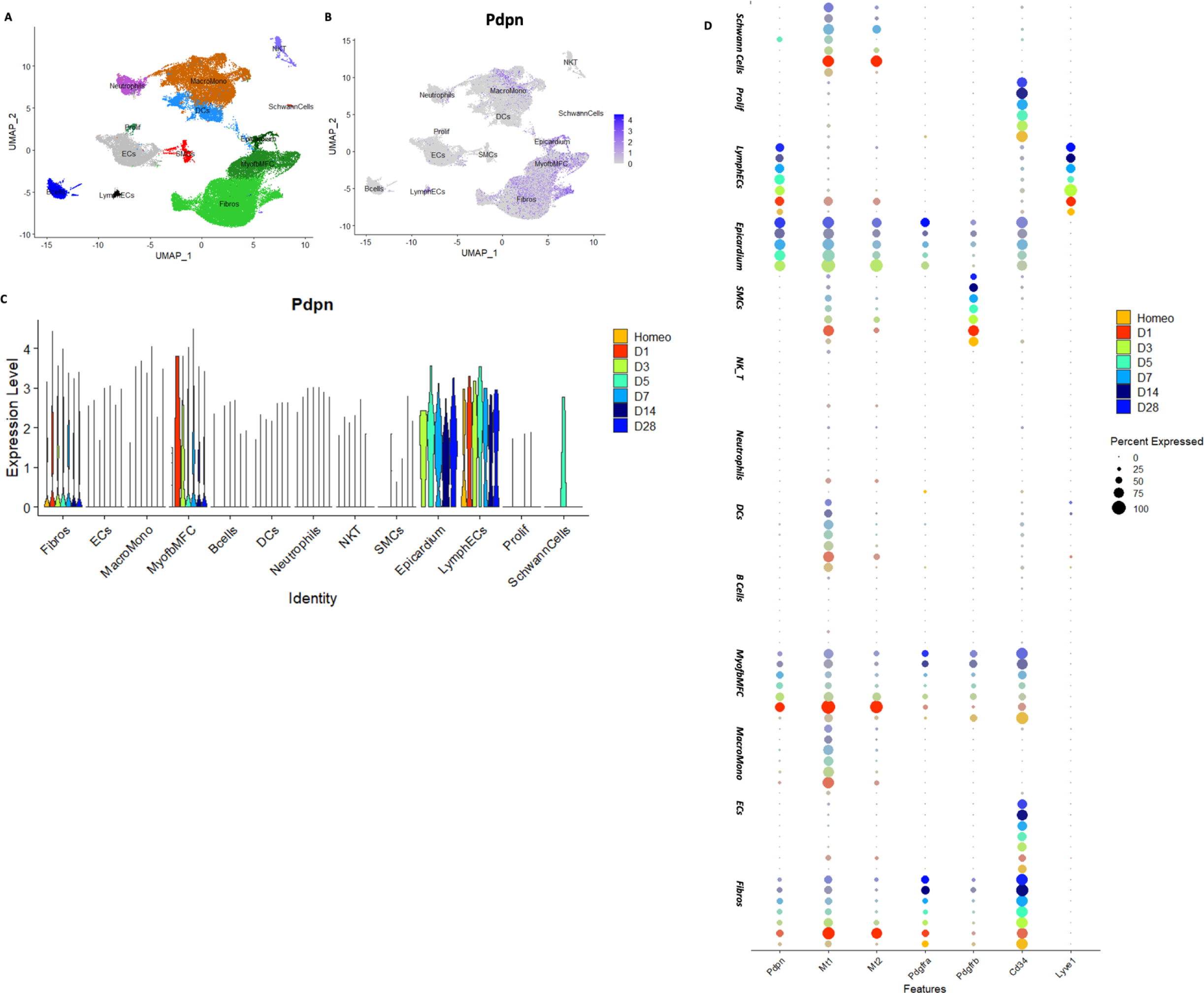
Single-cell RNA sequencing analysis identified Podoplanin (PDPN) expression in activated mesenchymal stromal cells (MSCs) post-myocardial infarction. (A) UMAP representation of the transcriptomic data from murine cardiac interstitial cells isolated by adult mouse cardiac ventricular tissue at homeostasis and 1,3,5,7,14,28 days post-ligation (reference to the papers) colored by cell population. (B) Feature plot indicating the expression intensity of PDPN. (C) Violin plot indicating the number of cells expressing PDPN per population, per time point. Fibros= Fibroblasts (3 subclusters), MonoMacro=Monocytes and Macrophages (4 subclusters), ECs = endothelial cells (2 subclusters), MyofbMFC= Myofibroblasts and Matrifibrocytes (2 subclusters), SMCs= smooth muscle cells, Prolif=Cells with high expression of proliferation genes. (D) Dot plot indicating the percentage of cell type expressing PDPN; Mt1, Mt2 (markers of injury response fibroblasts activated one-day post-MI (reference); PDGFRα, PDGFRβ, CD34 (markers of mesenchymal stromal cells) and Lyve1 (a marker of lymphatic ECs) at homeostasis, d1, d3, d5, d7, d14, d28 post-myocardial infarction. The dot size is proportional to the percentage of cells expressing the gene; the color indicates the time point and intensity of expression. This graphic representation confirms that activated mesenchymal stromal cells start the expression of PDPN soon after injury.

**Supplemental Figure 2.**
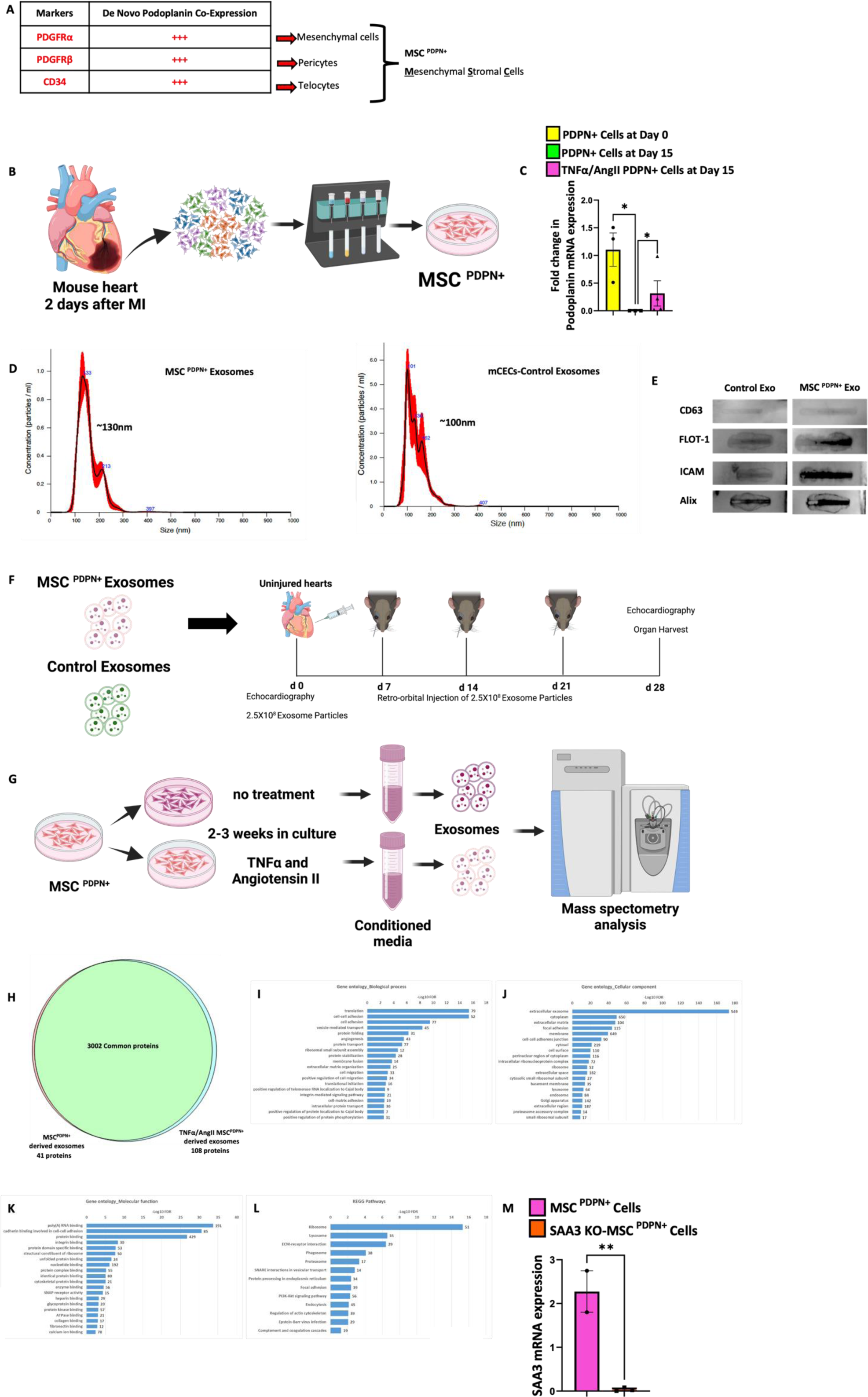
Related to Figures 1 and 2. Mesenchymal stromal cells are positive for Podoplanin (MSC^PDPN+^), and exosomes derived from MSC^PDPN+^ (MSC^PDPN+^Exosomes) can be successfully isolated in vitro and used for mass spectrometry analysis, in vitro and in vivo experiments. (A) In vivo, characterization of mesenchymal stromal cells positive for Podoplanin (PDPN) identified three main populations of cells that, as a sign of activation, acquire PDPN after ischemia. Mesenchymal cells positive for PDGFRα, pericytes positive for PDGFRβ, and telocytes positive for CD34 represent the heart’s most important cohort of mesenchymal stromal cells. (B) MSC^PDPN+^ were isolated two days after myocardial infarction (MI) from infarcted mouse hearts and cultured in vitro. (C) PDPN expression was reduced in culture conditions without stimulation and retained upon TNFα and Angiotensin II treatment. (D) Exosomes derived from pretreated MSC^PDPN+^ and control mouse cardiac endothelial cells (mCECs) were characterized for size using Nanosight and exosome protein marker array (E) to verify the quality of exosome isolation from cell-conditioned media. Data are presented as mean ± SEM, *P< 0.05. N=3-6. Ordinary one-way ANOVA analysis and Tukey’s post hoc test were performed among the groups. (F) 2.5×10^8^ Exosomes derived from pretreated MSC^PDPN^ or similarly treated mCECs were injected in the left ventricle of uninjured mouse hearts, followed by retro-orbital injection once every seven days. Mice were monitored with echocardiography and sacrificed 30 days after the first injection. Organs were harvested at the end of the study. (G) Exosomes, isolated from conditioned media of MSC^PDPN+,^ either pretreated or not with TNFα and Angiotensin II, underwent mass spectrometry analysis to investigate the protein content of the exosomal cargo. (H) Mass spectrometry analysis of untreated and TNFα and Angiotensin II pretreated MSC^PDPN+^ derived exosomes content revealed that ∼3000 proteins were commonly carried by the exosomes (and that only a few proteins were explicitly present in the untreated or TNFα and Angiotensin II treated group. (I) Specific gene ontology about the biological process, (J) cellular component, (K) molecular function, and (L) pathways activation confirmed that our samples are specifically extracellular vesicles. (M) q-PCR analysis showing that MSC^PDPN+^ isolated from SAA3 global knockout (KO) mice (SAA3 KO-MSC^PDPN+^) did not express mRNA for SAA3 when compared with wild-type (WT) MSC^PDPN+^. Data are presented as mean ± SEM, **P< 0.002. N=3. Student’s T-test analysis and Tukey’s post hoc test were performed among the groups.

**Supplemental Figure 3.**
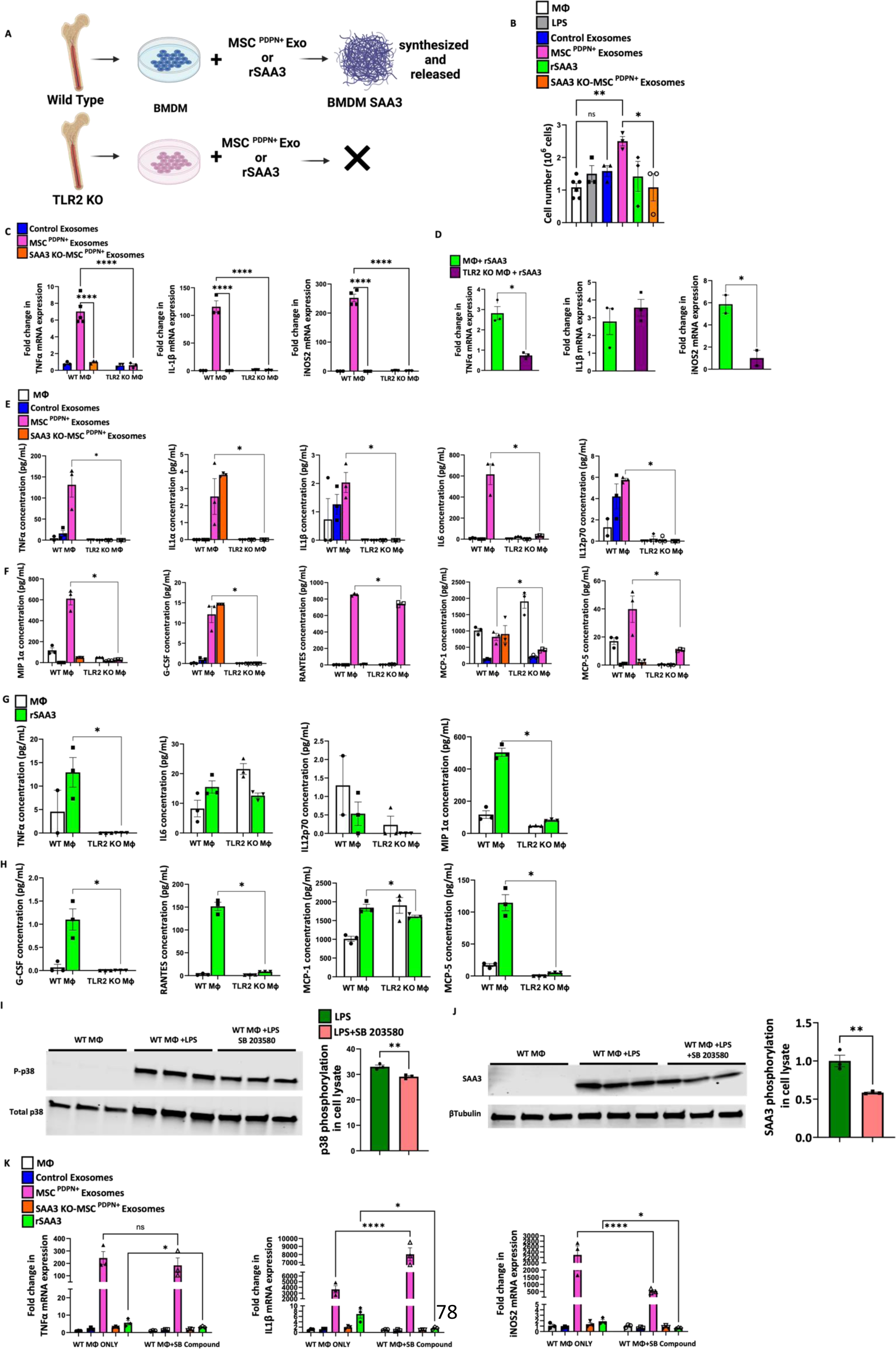
Related to Figure 4. Toll-like receptor 2 (TLR2) is required for bone marrow-derived macrophage activation when treated with recombinant Serum Amyloid a 3 (SAA3) and exosomes derived from mesenchymal stromal cells positive for Podoplanin (MSC^PDPN+^Exosome). (A) Bone marrow-derived macrophages (MΦ) isolated from wild-type (WT) and TLR2 knockout (KO) animals were isolated and cultured and treated either with exosomes derived from TNFα and Angiotensin II pretreated Podoplanin positive mesenchymal stromal cells (MSC^PDPN+^) or rSAA3. As shown in Fig. 4, TLR2 KO MΦ did not synthesize SAA3 upon stimuli. (B) Wild type MΦ migrated toward MSC^PDPN+^Exosomes and rSAA3 in a migration assay. (C) q-PCR analysis of wild type and TLR2 KO MΦ treated with exosomes derived from TNFα and Angiotensin II pretreated MSC^PDPN+^ and control exosomes (pretreated mouse cardiac endothelial cells-mCECs and SAA3 KO-MSC^PDPN+^ derived exosomes) and (D) q-PCR analysis of wild type and TLR2 KO MΦ treated with rSAA3 showed the expression of signature pro-inflammatory markers downstream of TLR2 activation. Data are presented as mean ± SEM, *P< 0.05, **P<0.002, ****P<0.0001. N=3-6. Ordinary one-way ANOVA or student’s T-test analysis and Tukey’s post hoc test were performed among the groups. (E) ELISA analysis of specific pro-inflammatory cytokines released by Wild type (WT) and TLR2 knockout (KO) bone marrow-derived macrophages (MΦ) treated either with exosomes derived from TNFα and Angiotensin II pretreated MSCPDPN+, control exosomes (pretreated mouse cardiac endothelial cells-mCECs derived exosomes), or SAA3 null pretreated MSCPDPN+Exosomes. (F) ELISA analysis of specific pro-inflammatory chemokines released by WT and TLR2KO MΦ after treatment with MSC^PDPN+^ Exosomes, control exosomes, or SAA3 KO-MSC^PDPN+^ Exosomes. Exosomes derived from SAA3 null pretreated MSCPDPN+ or MΦ null for TLR2 failed to be activated after treatments. (G) ELISA analysis of specific pro-inflammatory cytokines released by WT and TLR2KO MΦ after treatment with recombinant Serum Amyloid A 3 (rSAA3). (H) ELISA analysis of specific pro-inflammatory chemokines released by WT and TLR2KO MΦ after treatment with recombinant rSAA3. ELISA analysis showed enhanced expression of secreted cytokines and chemokines only in WT MΦ. MΦ null for TLR2 failed to be activated after rSAA3 treatment. Data are presented as mean ± SEM, *P< 0.05. N=3-6. Ordinary one-way ANOVA analysis and Tukey’s post hoc test were performed among the groups. (I, left) western blot analysis showing p38-MAPK phosphorylation in bone marrow-derived macrophages (MΦ) after treatment with lipopolysaccharides. Specific inhibition of p38-MAPK with SB 203580 compound reduced p38-MAPK phosphorylation in MΦ after treatment with LPS, quantified on the right. (J, left) Western blot analysis shows LPS-induced SAA3 synthesis in Mand consequent reduction of its expression after treatment with SB compound, with quantification on the right. (K) q-PCR analysis of MΦ after different types of treatments showed reduced TNFα, IL1β, and iNOS2 expression when treated with SB 203580 compound. Data are presented as mean ± SEM, *P< 0.05 and **P< 0.002. N=3. Student’s T-tests or ordinary one-way ANOVA analyses and Tukey’s post hoc tests were performed among the groups.

**Supplemental Figure 4.**
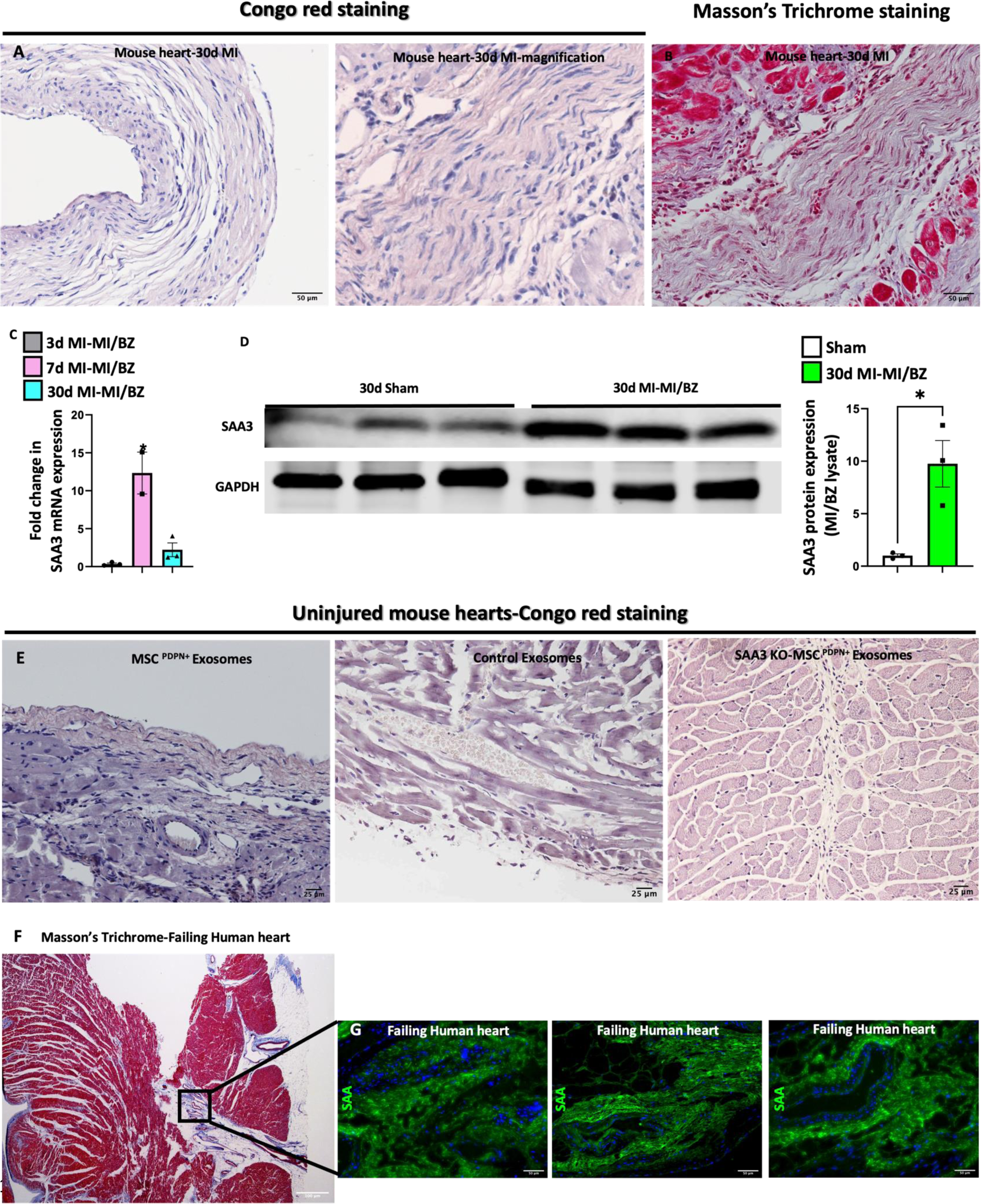
Related to Figure 5. Amyloidosis occurs in mouse hearts after myocardial infarction (MI) and in human failing hearts. (A) Congo red staining of mouse hearts after MI showed positivity for Congo red stain in the ischemic and fibrotic area (magnification in the middle) labeled in blue with Masson’s trichrome staining (B). (C) q-PCR analysis of the ischemic area of mouse hearts after MI showed that the peak of Serum Amyloid A 3 (SAA3) transcription corresponds to the inflammation phase when macrophages are activated. (D) Western blot analysis confirms the histochemistry data of SAA3 deposition in the ischemic area of infarcted mouse hearts after MI, with quantification on the right. Data are presented as mean ± SEM, *P< 0.05. N=3-10. Ordinary one-way ANOVA or student’s T-test analysis and Tukey’s post hoc test were performed among the groups. (E) Healthy mouse hearts treated with exosomes isolated from TNFα and Angiotensin II pretreated mesenchymal stromal cells positive for Podoplanin (MSC^PDPN+^) showed positivity to Congo red staining, left, when compared with healthy mouse hearts injected with exosomes isolated from pretreated mouse cardiac endothelial cells (mCECs), middle, or with healthy mouse hearts injected with exosomes isolated from pretreated MSC^PDPN+^ null for SAA3, right. (G) Extensive amyloid deposition in green has been observed in the same fibrotic area labeled in blue with Masson’s trichrome staining (F) in cardiac human sections of patients diagnosed with heart failure.

**Supplemental Figure 5.**
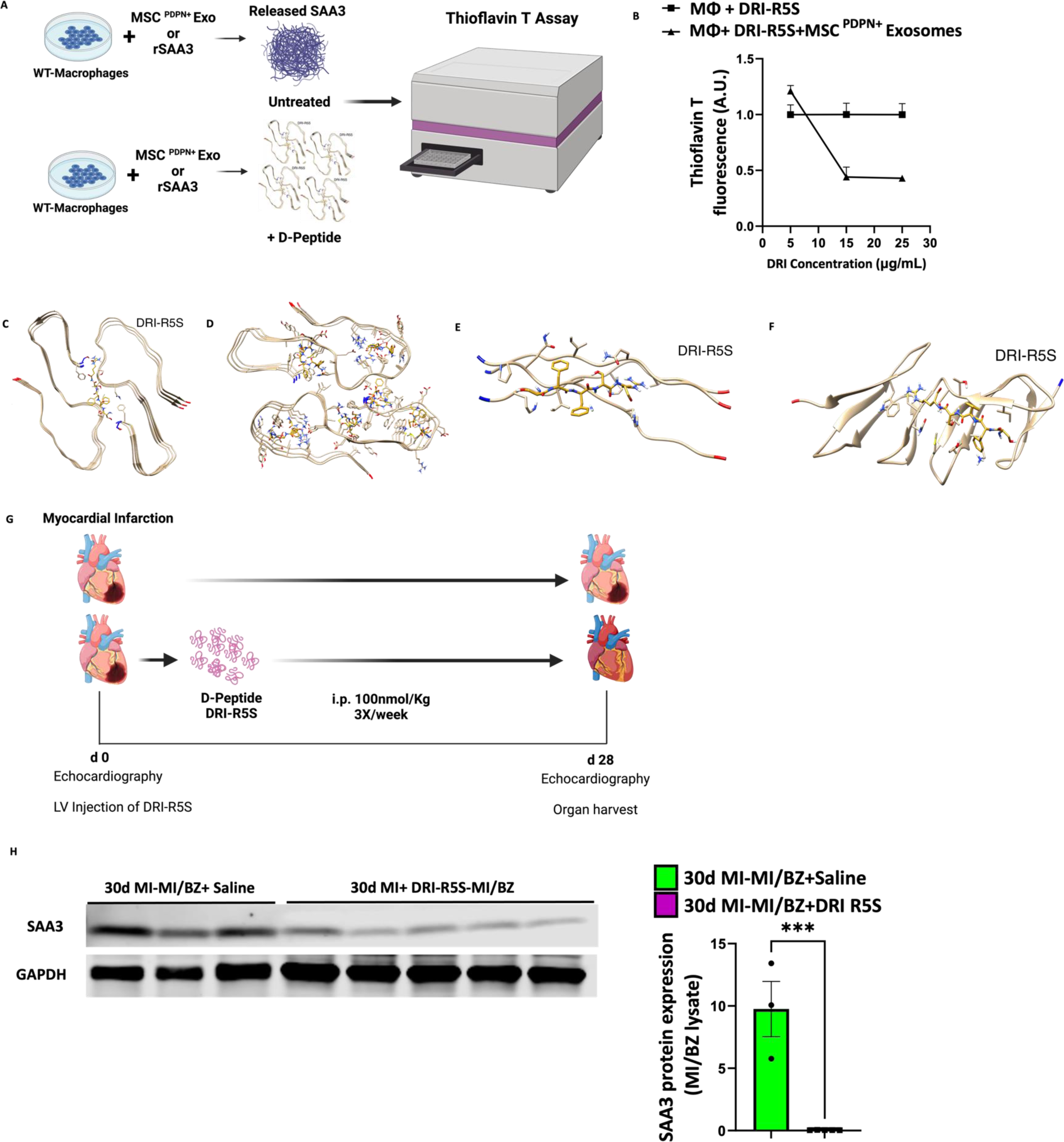
Related to Figures 6 and 7. D-Peptide DRI-R5S binds Serum Amyloid A 3 (SAA3) monomers when overexpressed by activated macrophages and treatment with DRI-R5S impaired Serum Amyloid A 3 (SAA3) aggregation and amyloid formation in mouse hearts after myocardial infarction (MI). (A) Wild-type (WT) bone marrow-derived macrophages (MΦ) treated either with exosomes derived from TNFα and Angiotensin II pretreated Podoplanin positive mesenchymal stromal cells (MSC^PDPN+^) or recombinant SAA3 (rSAA3), release SAA3 in the cultured media as shown in the ELISA of main Fig. 4, that can aggregate in amyloid structure. Using a D-Peptide DRI-R5S, we aimed to inhibit SAA3 aggregation. Thioflavin T assay measured SAA3 aggregates in the MΦ-conditioned media after treatment with and without DRI-R5S. (B) D-Peptide DRI-R5S docking with mouse SAA3 motif (main Fig. 7 A) reduced the aggregation of released SAA3 in M-conditioned media after treatment with exosomes isolated from TNFα and Angiotensin II pretreated mesenchymal stromal cells positive for Podoplanin (MSC^PDPN+^) in a dose-dependent manner. (C and D) DRI-R5S impairs the aggregation and destabilizes already formed Human SAA 1 cross-beta (β) fibers and fibril structures. (E) Collagen 1α and (F) fibronectin docked with DRI-R5S. (G) Wild-type (WT) animals underwent MI and were either treated or not with D-Peptide DRI-R5S once every other day for three weeks. (H) DRI-R5S treated animals showed reduced accumulation of SAA3 in the ischemic area, quantification on the right, compared to untreated animals injected with a DRI-R5S vehicle (saline). Data are presented as mean ± SEM, *P< 0.05, ***P<0.0005. N=3. Student’s T-Test analysis and Tukey’s post hoc test were performed among the groups.

**Supplemental Figure 6.**
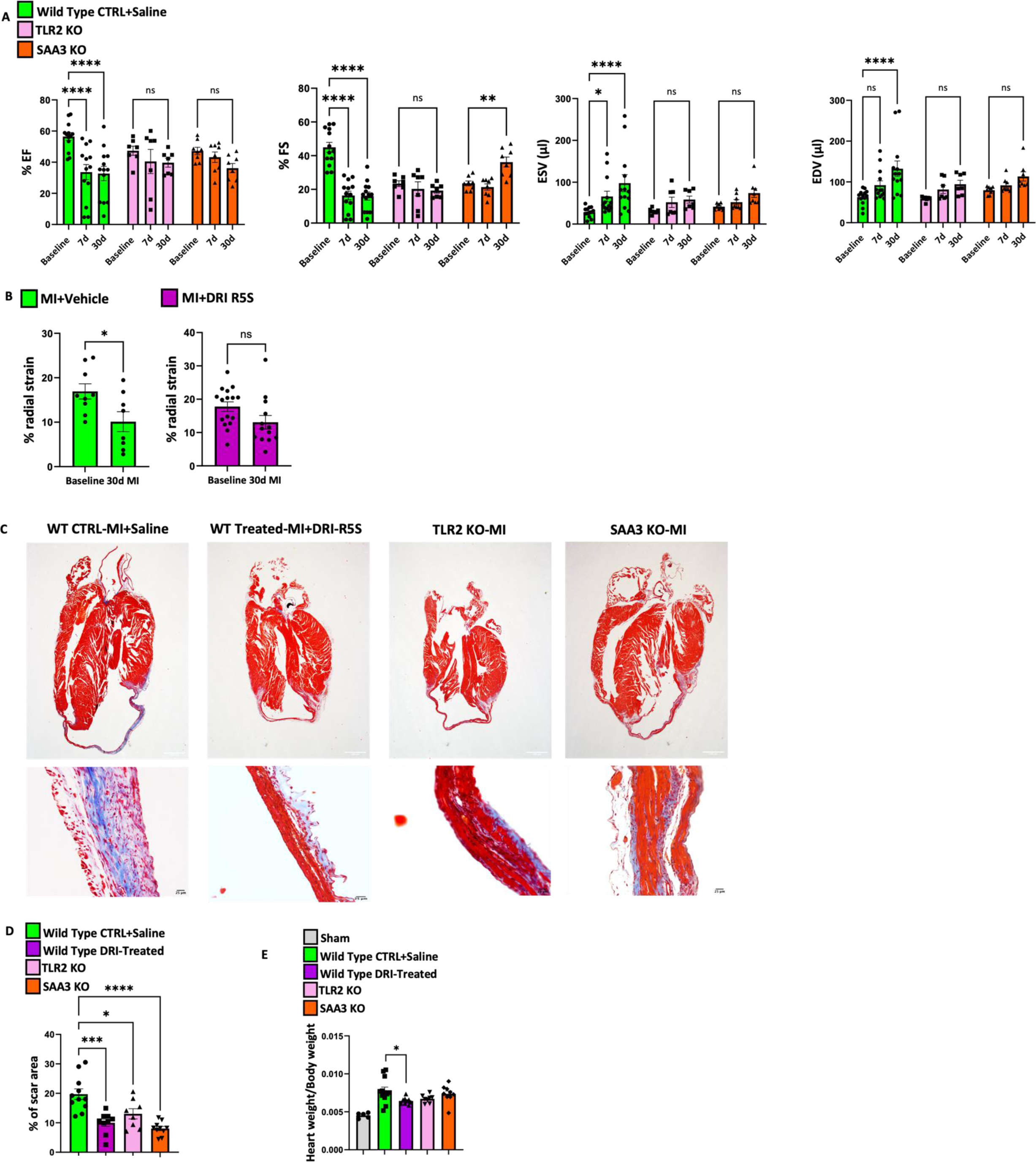
Related to Figure 7. Reduced Serum Amyloid A 3 (SAA3) amyloidosis leads to better heart function and scar formation post-myocardial infarction (MI). (A) Echocardiographic analysis of wild-type (WT), global Toll Like Receptor 2 (TLR2) knockout (KO), and global SAA3 KO animals that underwent MI for 30 days. TLR2KO and SAA3 KO animals showed improved cardiac function compared with wild-type animals after MI. Data are presented as mean ± SEM, *P< 0.05, **P<0.002, ****P<0.0001. N=7-10. Ordinary two-way ANOVA analysis and Tukey’s post hoc test were performed among the groups. (B) Analysis of % radial strain shows a statistical reduction in WT animals with MI and without any treatment (from 16.9% to 10.1 %). WT animals treated with DRI-R5S had a small reduction in % of radial strain, which wasn’t statistically significant (from 17.7% to 13.1%). Strain analysis shows an improved synchronous myocardial deformation potentially indicative of improved LV elasticity and, hence, improved LV contractility in mice hearts treated with DRI-R5S, although the presence of MI. Both groups of animals at baseline show similar % of radial strain (16.9% and 17.7%). Data are presented as mean ± SEM, *P< 0.05. N=7-10. Ordinary student’s T-test analysis and Tukey’s post hoc test were performed among the groups. (C) Histological analysis of wild-type animals either treated or not with D-Peptide DRI-R5S, TLR2, and SAA3 KO animals showed reduced cardiac scar size, quantified in D and better-left ventricle wall composition, magnification, in DRI-R5S treated and TLR2/SAA3 KO animals. (E) Quantification of heart weight/body weight ratio, data align with the quantified in C. Data are presented as mean ± SEM, *P< 0.05, ***P<0.0005, ****P<0.0005. N=7-10. Ordinary one-way ANOVA analysis and Tukey’s post hoc test were performed among the groups.

**Supplemental Figure 7.**
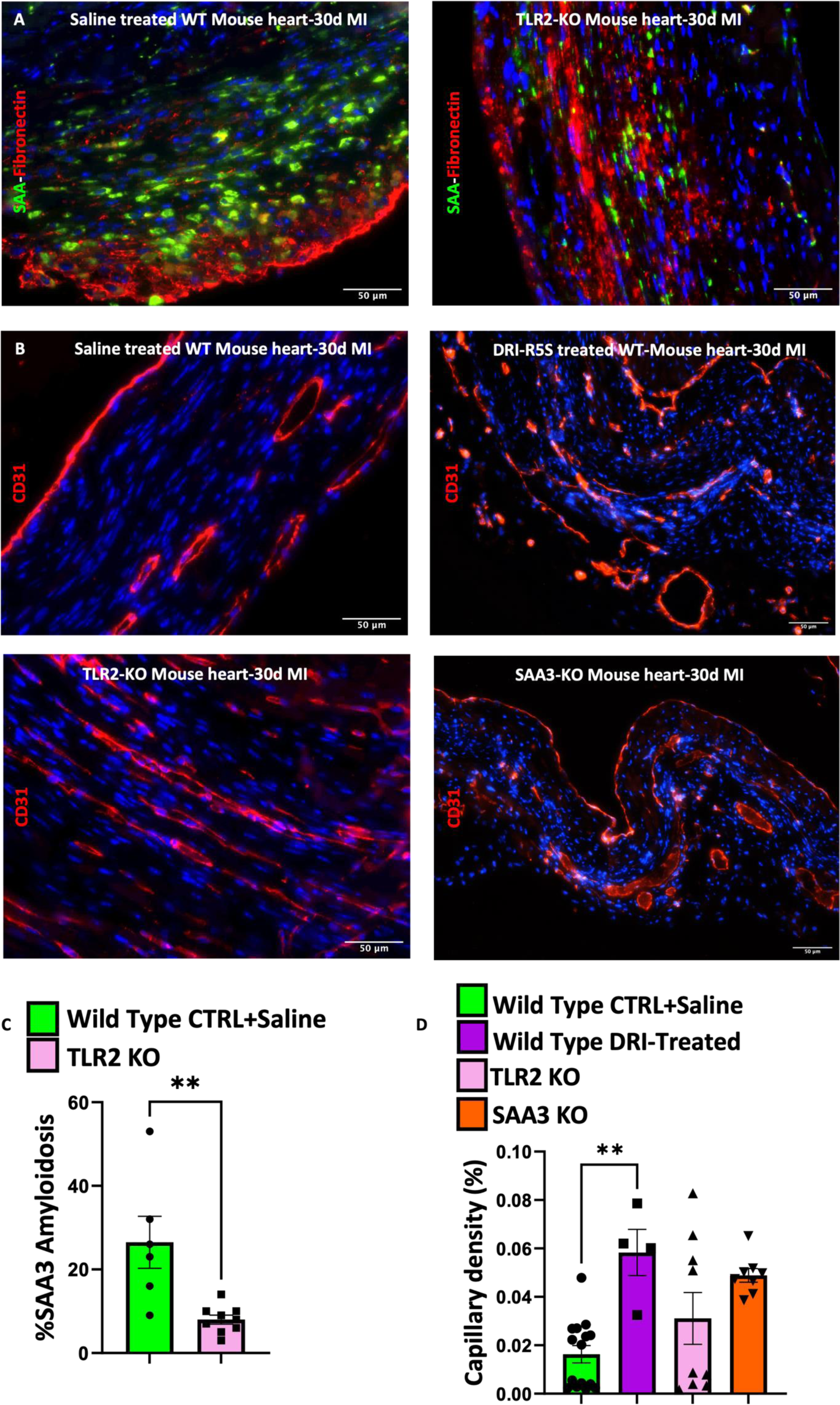
Related to Figure 7. Improved myocardial biogeography and vascularization are consequences of reduced Serum Amyloid A 3 (SAA3) amyloidosis. (A) Histological analysis of global Toll Like Receptor 2 (TLR2) knockout (KO) mouse hearts after myocardial infarction (MI) showed a reduction in SAA deposition, quantified in C when compared with wild-type mouse hearts 30 days after MI. Data are presented as mean ± SEM, **P< 0.002. N=7-10. Student’s T-test analysis and Tukey’s post hoc test were performed among the groups. (B) Histological analysis of wild-type (WT) animals either treated or not with D-Peptide DRI-R5S, TLR2KO, and global Serum Amyloid A 3 (SAA3) KO animals that underwent MI for 30 days. Protection from SAA3 amyloidosis improved myocardial scar vascularization in all animal groups compared to untreated animals. Blood vessels were stained with CD31 in red, and capillary density was quantified in D. Data are presented as mean ± SEM, **P< 0.002. N=7-10. Ordinary one-way ANOVA analysis and Tukey’s post hoc test were performed among the groups.

**Supplemental Table 1.** Literature search about Podoplanin acquisition by mesenchymal stromal and other cell types in different pathologies. The table groups all the pathologies that have been characterized by de-novo Podoplanin acquisition in mesenchymal stromal or other cell types. The type of cells that de-novo acquire Podoplanin are mostly mesenchymal stromal cells, unless the equivalent type of cells in the central nervous system or cancer tissue.

| Pathologies | Type of cells |
| --- | --- |
| Myocardial infarction and heart failure<sup>16-18,103,104</sup> | Mesenchymal stromal cells |
| Cardiac Fibrosis<sup>56,94,105</sup> | Mesenchymal stromal cells and early reactive fibroblasts |
| Thrombi and thrombo-inflammation<sup>97,106</sup> | Mesenchymal stromal cells |
| Deep vein thrombosis<sup>19,107</sup> | Mesenchymal stromal cells |
| Atherosclerotic lesion<sup>23,108-110</sup> | Mesenchymal stromal cells, Smooth muscle cells and macrophages |
| Aneurysm and High arteria shear<sup>85,111</sup> | Mesenchymal stromal cells |
| Calcification aortic valves<sup>112</sup> | Mesenchymal stromal cells and fibroblast |
| Stroke<sup>98</sup> | Microglia |
| Alzheimer Disease<sup>103</sup> | Mesenchymal and neural stromal cells |
| LPS-induced neuroinflammation<sup>113</sup> | Astrocytes and glial cells |
| Traumatic brain injury<sup>114</sup> | Astrocytes and glial cells |
| Gliosis and astrogliosis<sup>115</sup> | Astrocytes |
| Gliomas<sup>116,117</sup> | Astrocytes |
| Glioblastoma<sup>103</sup> | Mesenchymal and neural stromal cells |
| Gout arthritis, psoriatic arthritis, unclassified arthritis (UA), parvovirus associated arthritis, reactive arthritis<sup>118,119</sup> | Mesenchymal stromal cells and fibroblast |
| Rheumatoid arthritis<sup>118-123</sup> | Synovial fibroblasts |
| Psoriasis<sup>123,124</sup> | Synovial fibroblasts |
| Multiple sclerosis<sup>123</sup> | Mesenchymal stromal cells |
| Inflammatory bowel disease<sup>125</sup> | Mesenchymal stromal cells |
| Solid tumors<sup>126-128</sup> | Cancer cells and CAF |
| Cancer associated thrombi<sup>123,129</sup> | Cancer cells and Cancer associated fibroblasts |
| Intestinal segmented filamentous bacterial infection<sup>125</sup> | Epithelial cells |
| Hypercaloric diet<sup>130,131</sup> | Mesenchymal stromal cells |
| Cirrhosis<sup>132</sup> | Fibroblast |
| Preeclampsia<sup>133</sup> | Villous stromal cells |
| Fallopian tubes inflammation<sup>134</sup> | Telocytes |

**Supplemental Table 2.**
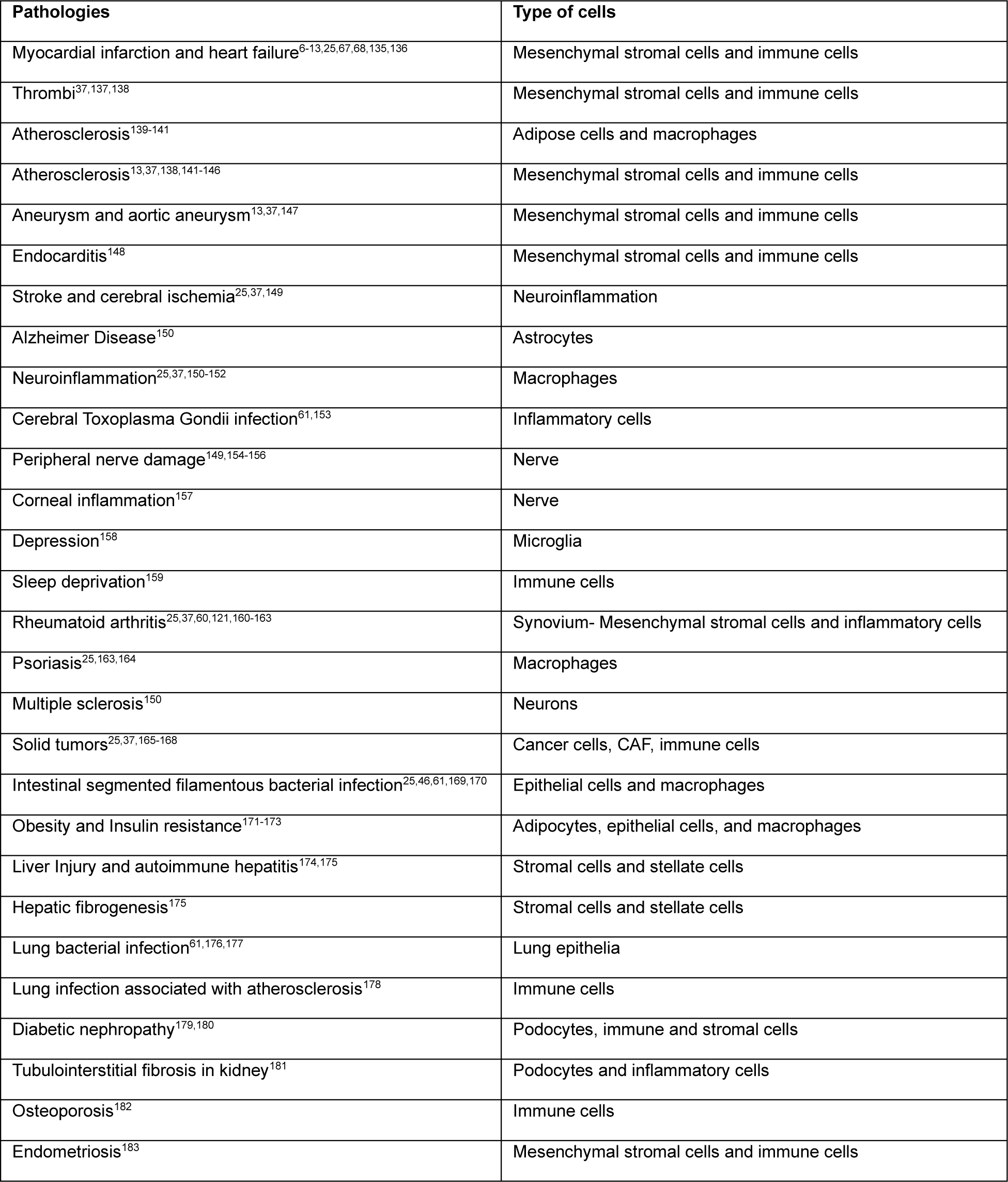
Literature search about Serum Amyloid A3 (SAA3) deposition as amyloidosis and acquisition by mesenchymal stromal and other cell types in different pathologies. The table groups all the pathologies that have been characterized by SAA3 deposition as amyloidosis and acquisition by mesenchymal stromal or other cell types. The type of pathologies in which SAA3 amyloidosis is present and SAA3 is the dominant cytokine are the same pathologies characterized by Podoplanin acquisition. Table 1 and table 2 correlate Podoplanin acquisition and SAA3 amyloidosis in different pathologies.

**Supplemental Table 3.** Antibodies used in the study.

| **Histological analysis antibody** | **Manufacturer** | **Catalog number** |
| --- | --- | --- |
| Fibronectin | Abcam | Ab2413 |
| CD45 | R&D | AF114 |
| SAA | Abbexa | Abx018140 |
| CD31 | R&D | AF3628 |
| Anti-Rabbit 555 | Invitrogen | A31572 |
| Anti-Goat 555 | Invitrogen | A21432 |
| Anti-Mouse 488 | Invitrogen | A21202 |

| **Western blot analysis antibody** | **Manufacturer** | **Catalog number** |
| --- | --- | --- |
| SAA3 | Abcam | Ab231680 |
| b-Tubulin | Proteintech | 66240-1 |
| GAPDH | Proteintech | 60004-1 |
| Phospho-p38 | Cell Signaling Technology | 28B10 |
| Total-p38 | Cell Signaling Technology | 9212 |
| Anti-Rat (680) | Li-Cor | 926-68076 |
| Anti-Mouse (800) | Li-Cor | 926-32212 |
| Anti-Rabbit (800) | Li-Cor | 926-32213 |
| Anti-Mouse (680) | Li-Cor | 926-68072 |

**Supplemental Table 3. Antibodies used in the study.**
| **FACS analysis antibody** | **Manufacturer** | **Catalog number** |
| --- | --- | --- |
| Podoplanin | R&D | BAF3244 |
| Lyve-1 | Abcam | Ab14917 |
| CD68 | Abcam | Ab31630 |

